# Allopurinol improves cardiovascular phenotypes of a mouse model for Williams–Beuren syndrome reducing redox stress

**DOI:** 10.1101/2025.08.06.668870

**Authors:** Alba Aizpuru-Gomez, Isaac Rodríguez-Rovira, Noura Abdalla, Jana Ruiz-Castro, Ana Paula Dantas, Gustavo Egea, Victoria Campuzano

**Affiliations:** Department of Biomedical Sciences, University of Barcelona School of Medicine and Health Sciences and IDIBAPS, 08036 Barcelona, Spain

## Abstract

Cardiovascular disease represents the primary cause of morbidity in Williams-Beuren syndrome (WBS), a neurodevelopmental disorder resulting from a hemizygous deletion of 26–28 genes on chromosome 7q11.23. The clinical phenotype includes systemic hypertension, cardiac dysfunction, and progressive vascular remodelling, features that are further exacerbated by increased oxidative stress. Given the early onset and progression of these cardiovascular manifestations, patients often require long-term pharmacological management from childhood. This raises important concerns about cumulative drug toxicity, dosing strategies, and the long-term safety of therapeutic interventions in pediatric populations.

Using the Complete deletion (CD) mouse model of WBS, this study aims to evaluate whether pediatric-equivalent doses of Allopurinol (ALO) (20mg/kg/day equivalent to 1,66mg/kg/day in humans) a highly specific xanthine oxidoreductase (XOR) inhibitor, and Losartan (LOS) (12,5 mg/kg/day equivalent to 1 mg/kg/day in humans) an angiotensin II type1 receptor antagonist, will exert beneficial effects on the cardiovascular phenotype of CD mice. To enable comparative analysis, the pharmacological effects of ALO and LOS were evaluated against both untreated CD mice and wild-type (WT) controls.

In CD mice, both treatments significantly reduced systemic blood pressure; however, their effects on cardiac outcomes were different. While treatment with ALO improved cardiac pathology, treatment with LOS did not. ALO likely acts by reducing XOR protein levels, reactive oxygen species (ROS) production and NRLP3 activation. These findings support a therapeutic model in which minimal doses of ALO offers broader cardiovascular protection in WBS.

## INTRODUCTION

Williams-Beuren syndrome (WBS) ([OMIM] 194050) is a rare genetic disorder attributed to a hemizygous microdeletion on chromosome 7q11.23 region. The microdeletion, which results in the loss of approximately 1.5 to 1.8 million DNA base pairs, is autosomal dominant and predominantly occurs de novo [1]. WBS patients present a constellation of distinct characteristics, including cardiovascular pathology, unique facial features, cognitive and developmental delays, and a distinctive behavioral profile characterized by hypersociability and anxiety-like behaviours [2].

Of the deleted genes, the *ELN* gene is of particular significance due to its involvement in elastin (ELN) synthesis. ELN in conjunction with fibrillin, forms the elastic fibers. Both are synthesized by vascular smooth muscle cells (VSMC) and are integral components of the extracellular matrix. Alongside collagen, which is produced by fibroblasts, they create a three-dimensional network that stabilizes the arterial structure. The deletion of one copy of *ELN* results in haploinsufficiency, which is crucial for explaining the cardiovascular phenotype associated with the syndrome [2]. Supravalvular aortic stenosis (SVAS) is a common and potentially life-threatening cardiovascular manifestation observed in individuals with WBS, often necessitating surgical intervention. Patients also present hypertension, hyperplasia of VSMC, connective tissue weakness and disorganized elastic fibers. The clinical presentation of WBS is multifaceted, with the severity and specific features varying among affected individuals due to factors such as the extent of the deletion and epigenetic modifiers[3,4].

Mechanistically, reactive oxygen or nitrogen species (ROS and RNS, respectively) levels have been pointed out as crucial mediators of cardiovascular phenotype development and progression in WBS. Hypertension and vascular stiffness were significantly less prevalent in patients whose deletion included *NCF1*, indicating that hemizygosity for *NCF1* was a protective factor [3,5]. Beyond this, mitochondrial dysfunction has emerged as an additional contributor to oxidative stress in WBS. This was first observed in mouse models, where altered mitochondrial structure and reduced ATP synthesis were associated with both cardiovascular and neurobehavioral phenotypes[6,7]. Notably, these mitochondrial alterations have also been confirmed in a neural biopsy from a WBS patient, supporting their translational relevance [6]. CD model exhibit persistent oxidative stress, which has been attributed to impaired *NRF2* signalling and upregulation of xanthine oxidoreductase (XOR) in cardiovascular tissues [8,9]. The resulting ROS contribute to endothelial dysfunction, smooth muscle cell proliferation, and inflammatory responses, definetively promoting SVAS and vascular stiffness [2,10].

Therapeutic approaches primarily focus on symptom alleviation. Thus antihypertensive agents offer symptomatic control given the high prevalence of hypertension, reported in a high percentage of affected children and even higher in adults [2]. Angiotensin receptors blockers (ARGs, Losartan/LOS) are attractive because they offer a dual benefit by not only controlling blood pressure but also by potentially ameliorating the processes that lead to vascular stiffness[4,11]. However, the long-term efficacy and safety of LOS therapy in WBS population remain under evaluation, particularly due to limited clinical and preclinical available data. We previous reported that some therapeutic effects on cardiovascular pathology were observed at high doses, there were also significant adverse outcomes[12]. This underscores the need for alternative therapeutic strategies that are both safer and more effective, ideally suitable for continuous administration from childhood ages (pediatric) through adulthood. In this context, allopurinol (ALO), a XOR inhibitor widely used in the treatment of hyperuricemia and gout [13], has emerged as a promising candidate for WBS therapy. ALO inhibits XOR, which is overexpressed in cardiovascular tissue [9], thereby reducing the production of intermediate metabolites such as ROS. The results show that ALO treatment improved cardiovascular features. This improvement could be correlated with the normalization of XOR expression and inflammasome signaling with the concomitant reduction of oxidative markers expression.

## MATERIALS AND METHODS

### Ethics statement

All procedures involving animals were performed in compliance with the National Institutes of Health’s Guide for the Care and Use of Laboratory Animals and approved by the local animal care committee of the Universitat de Barcelona (EB-310/22 and 192/22) in accordance with European (2010/63/EU) and Spanish (RD53/2013) regulations for the care and use of laboratory animals.

### Animals’ maintenance

CD mice, a WBS murine model that carries a 1.3 Mb heterozygous deletion spanning from *Gtf2i* to *Fkbp6*, were obtained as previously described [14]. All mice were maintained in a 97% C57BL/6J background. Genomic DNA was extracted from a mouse ear punch to perform genotyping using PCR and appropriate primers, as previously described [15]. Animals were housed under standard conditions in a 12 h dark/light cycle with access to food and water/treatment *ad libitum*. We used three groups of treatment (control, ALO, LOS) per genotype (WT and CD) in male and female mice, with n 7–11 in all groups.

### Treatment administration

Doses were established considering the pediatric range for both medications and in effective doses tested previously by the group [16]. The doses chosen are: 20mg/kg/day of ALO (equivalent to 1,66mg/kg/day in humans) (Sigma-Aldrich; CAS: 315-30-0) or 12 mg/kg/day of LOS (equivalent to 1 mg/kg/day) (Acros Organics; CAS: 124750-99-8). The equivalent human doses were calculated following Neur and Jacob (2016)[17]. Both compounds were prepared once every 2 days and dissolved in drinking water. LOS was maintained covered from light to prevent degradation. Respective treatments were started at 8–9 weeks old and maintained for 8 weeks until 16-17 weeks old.

The amount of drink per cage was quantified and normalized to the number of animals per cage (2–4) and to the time between each change (48–60h). Daily intake was not significantly different between groups (interaction time and treatment, F_(18,81)_= 1.652, *P*= 0.066 for males and F_(18,81)_= 0.559, *P*= 0.918 for females) (Supplementary Figure S1A). Both treatments significantly reduced body weight indepentdly of genotype (effect of treatment F_2,50_= 7.186, *P*=0.0018 in males; F_2,45_ = 22.97, *P*<0.0001 in females (Supplementary Figure S1B). This means that the animals are receiving an effective dose of ALO of 25-21 mg/kg/day for males and 26-25 mg/kg/day for females. For LOS we calculated effective doses of 14.6-12.6 mg/Kg/day for males and 17-14.2 mg/Kg/day for females. Supplemental Table S1 shows a detailed statistical analysis.

### Systolic Blood pressure measurement

The Systolic Blood Pressure (SBP) of conscious mice were measured using a tail-cuff system (Non-Invasive Blood Pressure System, PanLab) while holding mice in a retainer tube that was cleaned after each mouse. After habituation to the retainer, measures were taken on three separate occasions and averaged per mouse.

### Echocardiography

Two-dimensional transthoracic echocardiography was performed on all animals under 1.5% inhaled isoflurane. Imaging was conducted using a 10–13 MHz phased array linear transducer (IL12i, GE Healthcare, Madrid, Spain) with a Vivid Q system (GE Healthcare, Madrid, Spain). The images were recorded and later analysed offline using EchoPac software (v.08.1.6, GE Healthcare, Madrid, Spain). Proximal aortic segments were examined in a parasternal long-axis view, and the AR diameter was measured from inner edge to inner edge at end-diastole at the level of the sinus of Valsalva. The left ventricular end-diastolic (LVDD) and end-systolic (LVSD) internal diameters were measured in M-mode recordings and computed by the Teichholz formula [18]. The interventricular septum (IVSd) and left ventricular posterior wall thickness (LVWPd) in end diastole were also recorded. The average of three consecutive cardiac cycles was used for each measurement, with the operator being blinded to group assignment.

### Sample obtention

For histological analysis, tissues were fixed in 4% paraformaldehyde and embedded in paraffin. For protein analysis, samples were dry-frozen and stored at −80 °C.

Hearts and aortas measurements were carried out in a blinded manner by two different observers with no knowledge of genotype and treatment. Images were captured using a Leica Leitz DMRB microscope (10x objective) equipped with a Leica DC500 camera and analysed with Fiji Image J Analysis software.

### Western Blot

Dissected tissues were homogenized in RIPA buffer with protease (PMSF, aprotinin, leupeptin, pepstatin) and phosphatase (Na₃VO₄, NaF) inhibitors using a bullet blender and beads. Protein concentration was measured with the DC Protein Assay Kit (Bio-Rad). Membranes were incubated overnight at 4 °C with primary antibody against XOR (1:500, Santa Cruz sc-398548) and NRLP3 (1:500, Sigma-Aldrich ZRB1762). After washing, membranes were incubated with HRP-conjugated secondary antibody (1:3000, anti-rabbit, Promega W401B) and visualized using Luminol reagent (Santa Cruz). Bands were quantified using ImageJ (v1.53).

### Immunohistochemistry and inmunohistofluorescence staining

Paraffin-embedded heart TMAs (5 μm) were deparaffinized, rehydrated, and subjected to antigen retrieval in 10 mM sodium citrate buffer (0.05% Tween, pH 6) for 30 min at 95 °C in a steamer. Free aldehyde groups were incubated with 50 mM NH₄Cl (pH 7.4) for 20 min, followed by permeabilization with 0.3% Triton X-100 for 10 min. Horseradish peroxidase (HRP)-based immunohistochemistry was used to stained aortic tissue sections for 3-‘NT and XOR. After PBS washes, sections were blocked with 1% BSA in PBS for 2 h at room temperature and incubated overnight at 4 °C with anti-3-NT (1:200; Merck Millipore, 06-284) or XOR (1:50; Rockland 200-4183S). On the next day, sections were incubated with goat anti-rabit secondary solution (1:500; abcam ab97051) for 1 hour followed by the liquid DAB+Substrate Chromogen system (K4065) for 1 minute at room temperature or 1 hour with Alexa Fluor 647-conjugated goat anti-rabbit secondary antibody (1:1000; Invitrogen, A-21246), counterstained with DAPI (1:10,000) and imaged using an AF6000 Widefield fluorescent microscope. HRP-stained sections were visualized under a Leica Leitz DMRB microscope at 40X. Quantification was performed on four regions per section using ImageJ (v1.53).

### Sirius Red Staining

Paraffin-embedded heart and aorta sections (4 μm) were deparaffinized, incubated in 0.2% phosphomolybdic acid for 2 minutes, and stained with Sirius Red in saturated picric acid for 110 minutes in a humid chamber. Sections were washed, treated with 0.01 N HCl, briefly destained in 75% ethanol, dehydrated, cleared in xylene, and mounted with DPX. Quantification was performed on four regions per section using ImageJ (v1.53).

### Statistical analysis

Data are presented as mean ± standard error of the mean (SEM). Normality and homogeneity of variances were assessed using the Shapiro–Wilk tests, and Levene’s test, respectively. Group differences were analysed by one-way or two-way ANOVA followed by Tukey’s post hoc test when assumptions were met, or by Kruskal–Wallis with Dunn’s post hoc test otherwise. Outliers were identified and excluded using the ROUT method (Q = 1%). Values were considered significant when p < 0.05. GraphPad Prism 10 software (version 10.0.0) was used for obtaining all statistical tests and graphs.

## RESULTS

### LOS and ALO treatments normalize systolic blood pressure of CD mice

As previously reported [19], CD mice exhibit elevated systolic blood pressure (SBP) compared to their wild-type (WT) littermates. To validate the selected dosing regimen, SBP was measured as an outcome.

At 2-month-old CD mice shown a significant increase in SBP (T_(56)_= 7.238; *P*<0.0001; Figure 1A). Consistent with the hypertensive phenotype characteristic of CD model at 4 months of age, SBP was significantly higher in untreated CD mice. Two-way ANOVA revealed a significant factor interaction genotype/tratment (F_2, 117_= 46.91; *P*<0.0001). Pair comparisons shown that both treatments significantly reduced the characteristic high SBP occurring in CD mice (P<0.0001), the values obtained being no significantly different to WT animals (P= 0.5678 for ALO and P= 0.8982 for LOS; Figure 1B). No significant differences were found between the two treatment groups, nor between the WT and treated control groups (*P*=0.5095 for ALO and *P*=0.6093 for LOS), averaging around 115 mmHg. Results crearly underline the effectiveness of LOS and ALO in the normalization of SBP in the CD model. See Suplemental Table S2 for detaliled statistical analysis.

**FIGURE 1:**
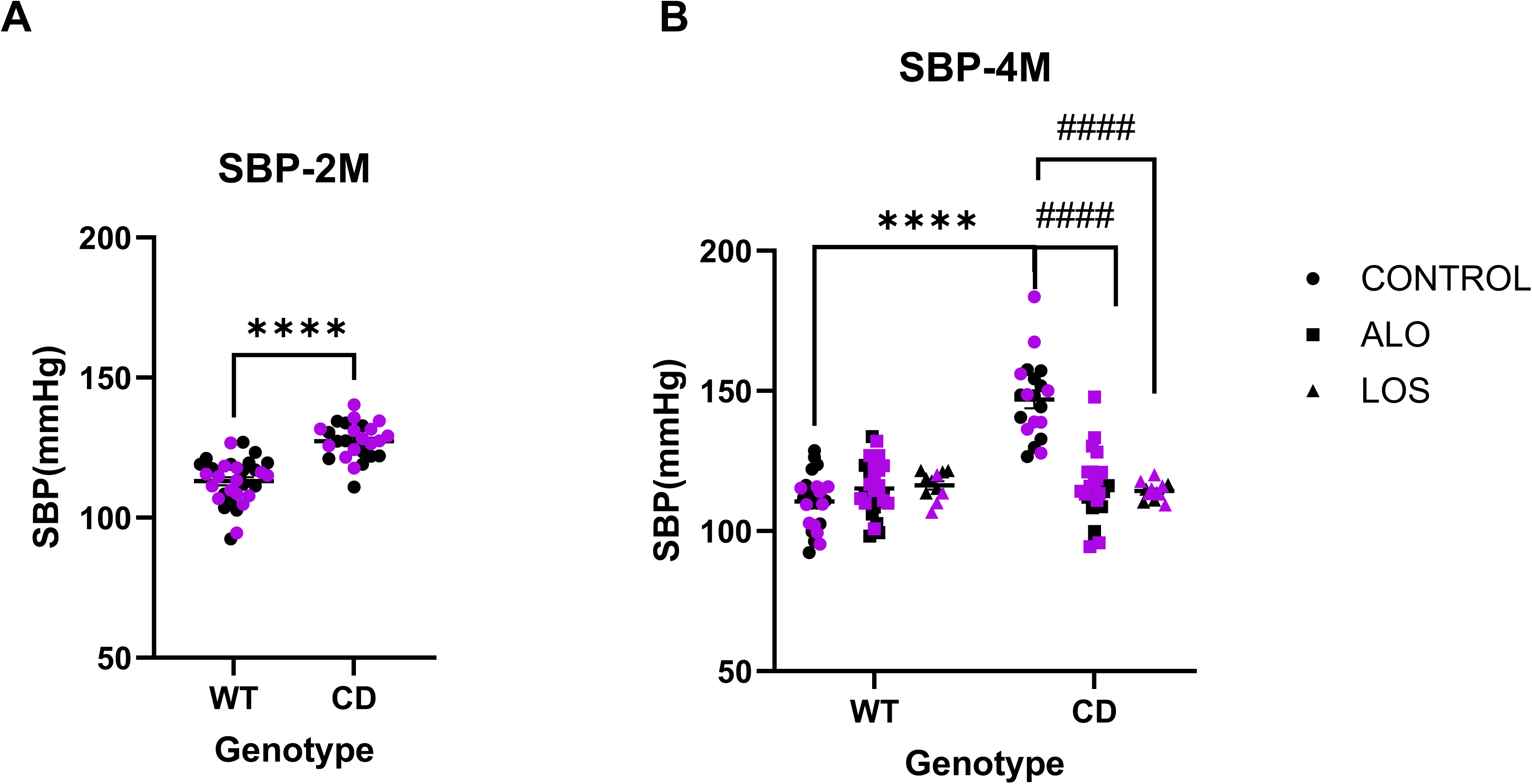
Systolic blood pressure normalization in treated CD mice**. (A)** Basal levels of SBP in the conscious 2-month-old mice. Statistical significance was calculated by Student’s T test**. (B)** SBP measured WT and CD conscious mice treated for 8 weeks with ALO (20 mg/kg/day), LOS (12,5mg/kg/day) or untreated (control). Statistical significance was calculated by Tukey’s *post hoc* test following two-way ANOVA. Note the significant differences (*P*<0.0001) between treated and untreated CD animals. There are no significant differences between the treated CD animal groups (P=0.9997). Circles, Control; Squares, ALO treatment; Triangles, LOS treatment. Black, males; Purple, females. *Genotype effect; ^#^ treatment effect. ****^,####^ p<0.0001. Data are expressed as mean ± S.E.M

### LOS and ALO treatments improve CD-associated ascending aortic stenosis

Next, we evaluated the effect of LOS and ALO treatments on the characteristic ascending aortic stenosis present in the CD model. We first evaluated, using echocardiography, the status of the stenosis before the start of treatment (2 months of age). A significant decrease in aortic root diameter was observed in the CD animals (Figure 2A).

**FIGURE 2.**
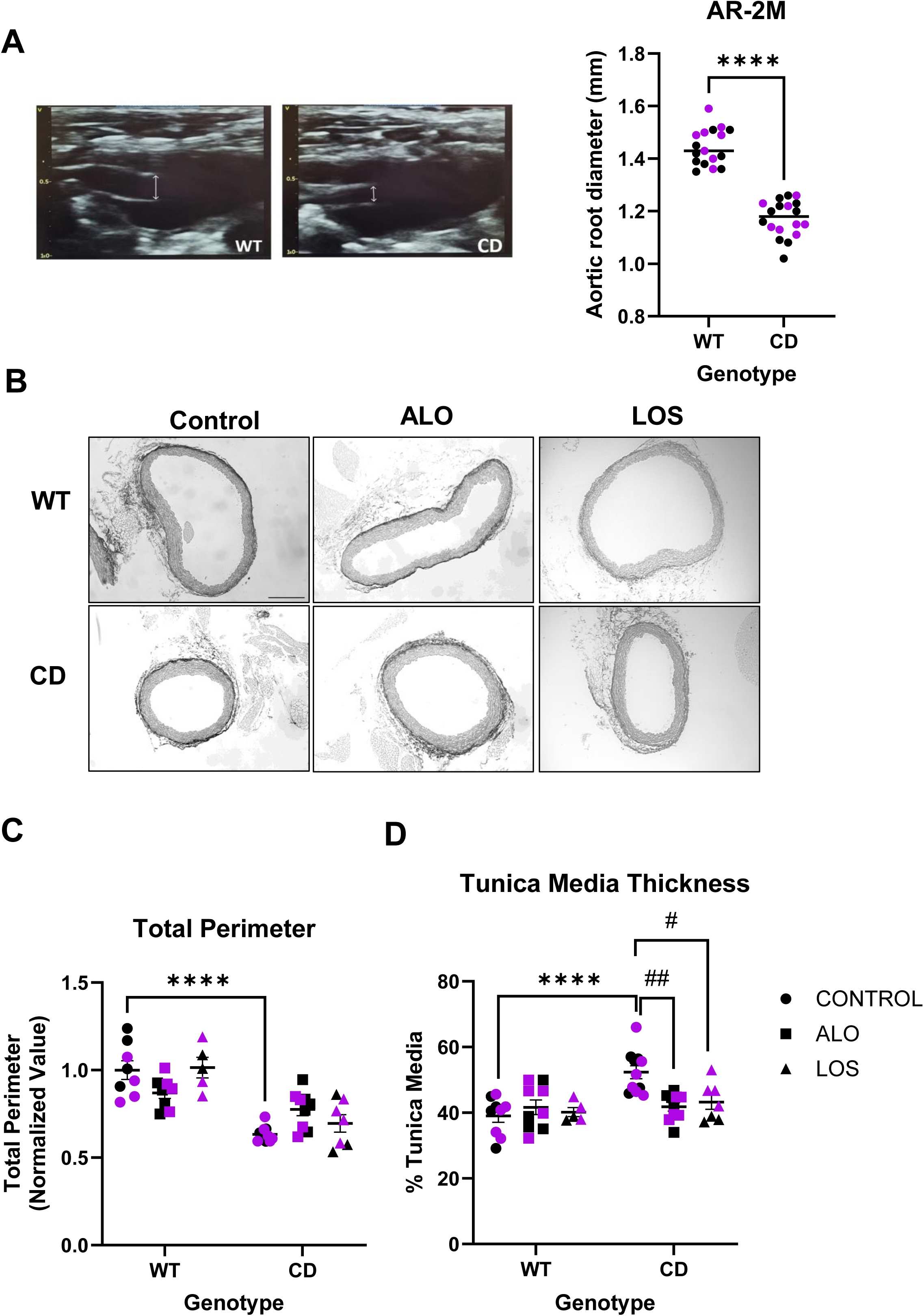
Treatment improves aortic stenosis-associated to CD mice. **(A)** Stenosis is already present in CD mice at 2-month-old. (Left), Representative images of aortic root obtained by echocardiography. (Right), Histogram representing the cuantification of aortic root diameter. Values normalized to WT. Statistical significance was calculated by Student’s T test. **(B)** Representative images of ascending aortas slides stained with Picrosirius Red. 10X magnification. Scale bar, 500 µm. **(C)** Histogram representing the total diameter quantification of ascending aorta and **(D)** Thickness of the tunica media represented as a percentage of the total area of the aorta Values normalized to WT-Control.Statistical significance was calculated by Tukey’s *post hoc* test following two-way ANOVA. Circles, Control; Squares, ALO treatment; Triangles, LOS treatment. Black, males; Purple, females. *Genotype effect; # treatment effect. ^#^ *P*<0.05; ^##^ P<0.01; *** P<0.001; **** P<0.0001. Data are expressed as mean ± S.E.M.

After two month of treatment (4 month of age), we evaluated treatment effects in aortic wall organization by histomorphology, quantifying the total perimeter and thickness of the tunica media in the ascending aortas as previously defined [9]. Histological analysis revealed that untreated CD mice continued to exhibit significantly decreased aòrtic perimeter and increased aortic wall thickness compared to WT controls (P<0.0001; Figure 2 B,C and D), consistent with the vascular remodelling associated with SVAS. Neither LOS nor ALO significantly affected the aortic perimeter, which remained significantly smaller (P=0.0016 for ALO and P<0.0001for LOS; Figure 2C). However, CD-treated mice significantly reduced aortic wall thickness compared to untreated-CD mice (P=0.0014 for ALO, and P=0.0205 for LOS; Figure 2C). Notably, there were no significant differences between the CD-treated groups and WT control, indicating full normalization of this parameter. This reduction in the thickness of the tunica media is probably due to a reduction in cell density (treatment effect F_(2,34)_=9.986, P=0.0004; Figure S2). All statistical analysis was detailed in Table S3.

### ALO is more effective than LOS improving cardiac function in CD animals

To evaluate the effect of both treatments on the cardiovascular phenotype of CD mice several anatomical and functional parameters were evaluated by transthoracic echocardiography.

First, we evaluated cardiac dysfunction through ejection fraction (EF) before starting any treatment (2 month old mice). As expected, CD mice presented a reduction in EF compare to WT littermates (P=0.003; Figure S3).

We hypothesized that the observed improvement of SBP and aortic stenosis by both treatments should improve the cardiac function. The analysis after two months treatment revealed that CD mice presented a reduction in left ventricular ejection fraction (EF) both in males and females compare to WT littermates (effect of genotype, P<0,0001; Figure 3A). A significant effect of threatment could be appreciated in both males and females (effect of treatment, P<0,0001 for males and P=0.0119 for females). Results indicated that treatments produced an overall improvement in cardiac function evidenciated by the normalization of the EF in treated CD mice.

**FIGURE 3:**
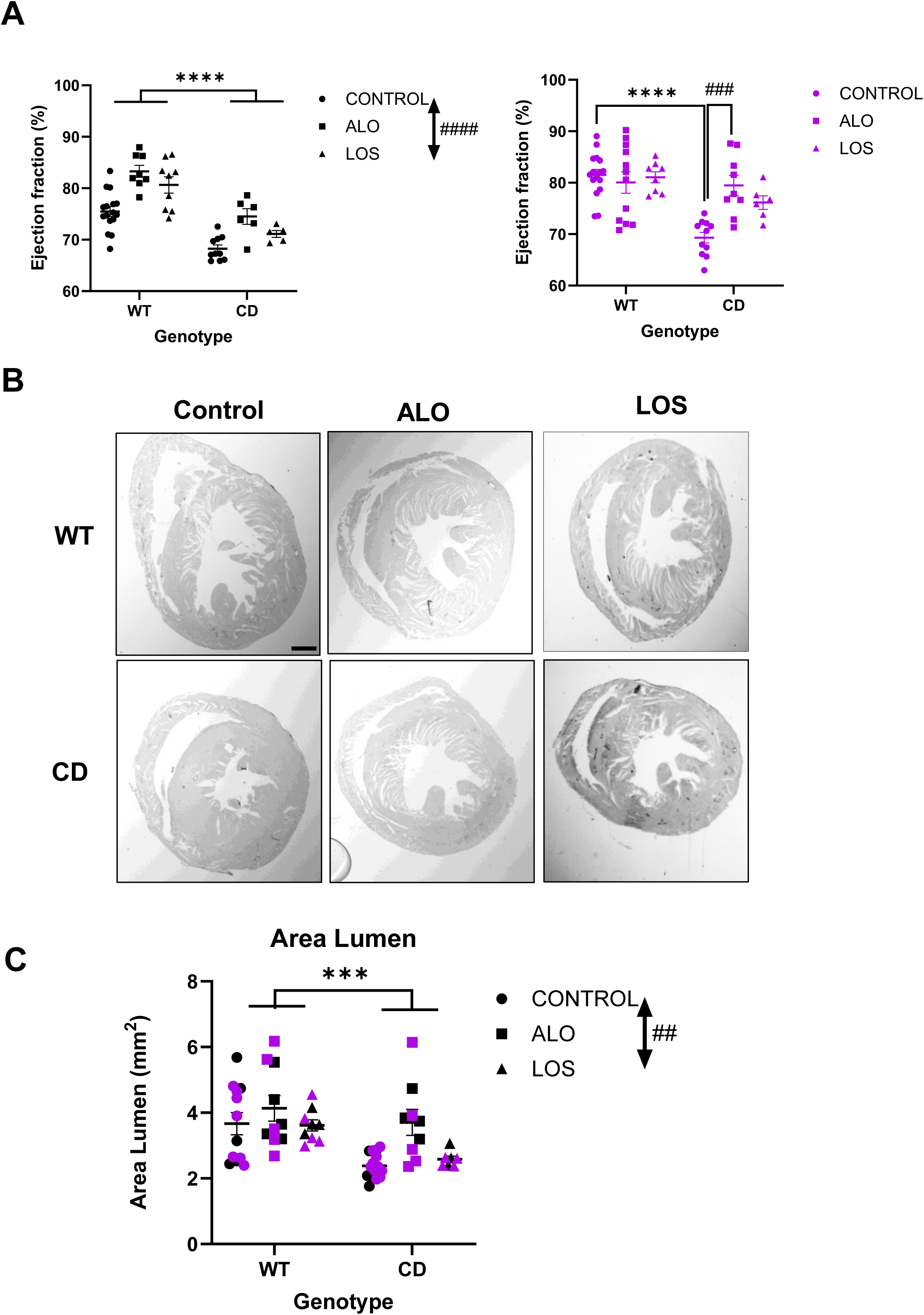
Cardiac dysfunction-associated to CD mice improves with ALO treatment. **(A)** Ejection Fraction increased after treatment, both in males (left) and females (right). Note the significant different between ALO-treated and untreated-CD females (*P*=0.0004). Statistical significance was calculated by two-way ANOVA followed by Tukey’s *post hoc* test in females. **(B)** Representative images of transversal heart sections stained with Picrosirius Red. 4X magnification. Scale bar, 500 µm. **(C)** Histogram representing the quantification of lumen area. Statistical significance was calculated by two-way ANOVA. Circles, Control; Squares, ALO treatment; Triangles, LOS treatment. Black, males; Purple, females. *Genotype effect; # treatment effect. ^##^ P<0.01; ***^,###^ P<0.001; **** P<0.0001. Data are expressed as mean ± S.E.M.

To further validate these changes, the area of the left ventricular lumen was evaluated by histological analysis. In accordance with echocardiographic observations, treatment significantly increased the area of the left ventricular lumen (effect of treatment, *P*=0.0043, Figure 3B). All statistical analysis was detailed in Table S4.

### ALO but not LOS reduces cardiac hypertrophy of CD mice

We also evaluated echocardiographic parameters related to hypertrophy since this phenotypic trait is characteristic of CD animals. Wall thickness at the interventricular septum (IVSd) and the posterior wall (LVPWd) were significantly increased in CD mice both in males and females. After LOS and ALO treatments there were a trend towards a decrease in IVSd which became only significant in females (treatment effect, F_2, 58_= 3.442; *P*= 0.0387) (Figure 4A and Figura S4).

**FIGURE 4:**
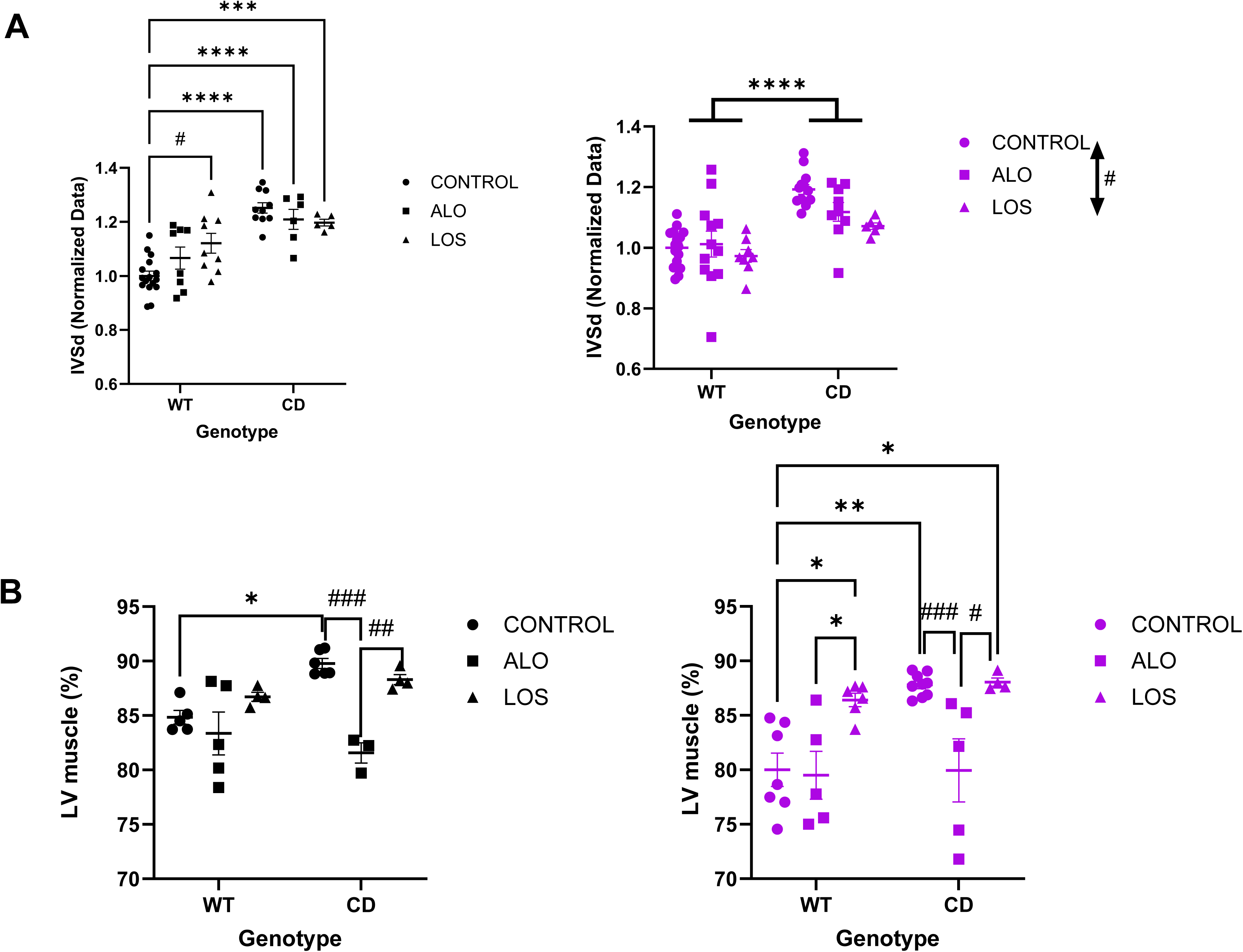
Heart hypertrophy present in CD mice improves with ALO treatment. (**A**) Interventricular septum thickness in diastole (IVSd) measured by echocardiography. showed a significant thickening of the IVSd both in males and females (genotype effect *P*<0.0001) with significant reduction after treatment in females (treatment effect, *P*=0.0387). Statistical significance was calculated by two-way ANOVA followed by Tukey’s *post hoc* test in males. (**B**) Thickness of the left ventricular muscle represented as a percentage of the total area of left ventricle. The significantly increased thickening of the left ventricular wall identified in untreated CD mice (*P*=0.0144 for males and P= 0.002 for females, genotype effect) was normalised only following ALO treatment (*P*=0.0004 for males and *P*= 0.0055 for females, treatment effect). Circles, Control; Squares, ALO treatment; Triangles, LOS treatment. Black, males; Purple, females. *Genotype effect; ^#^ treatment effect. *^,#^ *P*<0.05; **^,##^ *P*<0.01; ***^,###^ *P* < 0.001; **** *P* < 0.0001.

Histologicaly, cardiac hypertrophy was assessed by measuring the muscle proportion of the left ventricle. Consistent with the presence of hypercardiopathy in untrated CD animals, we observed a significant increase in muscle percentage, both in males and females (P=0.0144 and P= 0.002, respectively, Figure 4B). Interestingly, ALO but not LOS, significantly reduced the muscle percentage of left ventricle in CD animals, reaching values similar to those of untreated WT controls (P = 0.3535 in males and P> 0.999 in females). All statistical analysis were detailed in Table S5.

### ALO reverses XOR overexpression and 3’NT levels in cardiovascular tissues of CD mice

We were next interested in uncovering the mechanisms by which ALO might contribute to explaining the increased benefits of cardiac phenotypes associated with WBS. The severity of the cardiovascular phenotype in WBS is highly dependent on oxidative stress, which among other factors, could be influenced by XOR overexpression [9]. One of the mechanism by which ALO could contribute to these improvement is regulating pathways and processes related to oxidative stress [20,21].

First, we analyzed the effect of ALO treatment in XOR expression. We observed a significant increase in basal levels of XOR in myocardium (Figure 5A) of CD mice (genotype effect, *P*<0.0001). ALO treatment significantly attenuated this overexpression (treatment effect, *P*< 0.0001).

**FIGURE 5:**
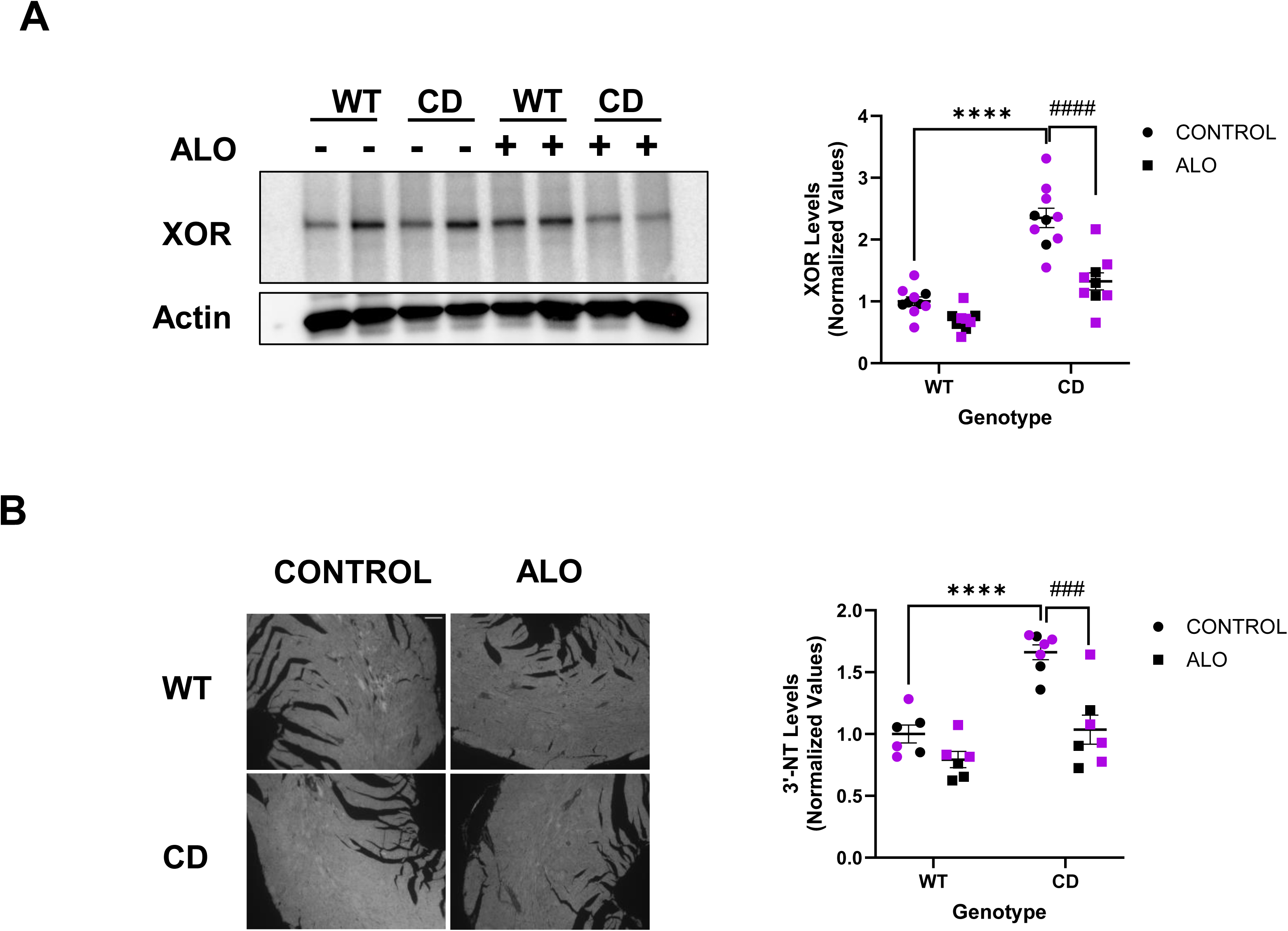
ALO treatment reduces XOR and 3NT levels in myocardium of CD mice. **(A)** In myocardium XOR expression was analysed by western blot (left). Quantitative analysis showed a significant reduction in ALO-treated versus untreated CD mice (*P*<0.0001)**. (B)** Representative images of 3NT immunofluorescence detection in myocardium (left). Scale bar: 200 µm. We observed a significant genotype/treatment interaction F_(1, 22)_ = 6.084 *P*= 0.0219) for 3-NT levels. Values normalized to WT-Control. Circles, Control; Squares, ALO treatment. Black, males; Purple, females. Statistical significance was calculated by Tukey’s *post hoc* test following two-way ANOVA. *Genotype effect; ^#^ treatment effect. ^###^*P* < 0.001; ****^,####^*P* < 0.0001.Data are expressed as mean ± S.E.M.

Second, to determine whether the reduction in XOR levels was concomitant with a reduction in oxidative stress, we analyzed the expression levels of the 3’NT marker (Figure 5B). A two-way ANOVA showed that the significantly elevated levels present in the myocardium of CD model (genotype effect, *P*< 0.0001) were attenuated after ALO treatment (treatment effect, *P*< 0.0001) (Figure 5B). Similar results were obtained for ascending aortae, both in XOR overexpression (Figure S5A) and 3NT levels (Figure S5B), which pointed towards a potential protective effect of ALO reducing oxidative damage in cardiovascular tissue.

All statistical analysis were detailed in Table S6.

### ALO attenuates NLRP3 inflammasome activation of CD mice

Another complementary mechanism by which ALO could contribute to explaining the beneficts in WBS-cardiac phenotype is regulating processes related to inflammation [22–24]. It has been demonstrated that XOR inhibition attenuates nucleotide-binding domain, leucine-rich containing family, pyrin domain-containing 3 (NLRP3) inflammasome activation [25]. In addition, pressure overload to the left ventricle activates NLRP3 inflammasome in cardiomyocytes, fibroblasts, and vascular endothelial cells and contributes to cardiac hypertrophy [26]. We had previously described an increase in NLRP3 inflammasome activation in brains of CD mice [27], but unkown in CD hearts.

We therefore analysed NLRP3 levels in myocardium of CD mice. We observed a significant increase in basal levels of NLRP3 in CD mice compared with WT littermates (genotype effect, *P* <0.0001). After ALO treatment, NLRP3 levels were normalised (*p*=0.8817) (Figure 6). All statistical analysis were detailed in Table S7.

**FIGURE 6:**
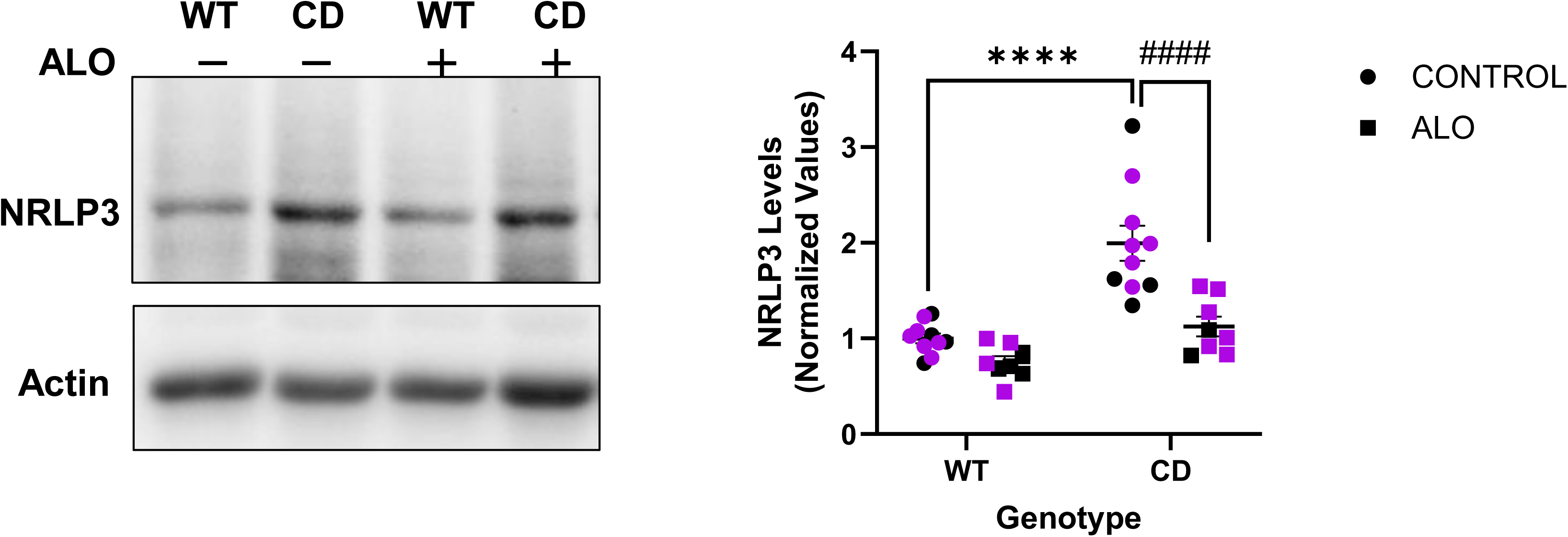
ALO treatment reduces NRLP3 levels in myocardium of CD mice. NRLP3 expression levels analysed by western blot (left) in myocardium and the corresponding quantification (right). Note the significant differences between untreated and ALO treated CD mice (*P*<0.0001). Values normalized to WT-Control. Circles, Control; Squares, ALO treatment. Black, males; Purple, females. Statistical significance was calculated by Tukey’s *post hoc* test following two-way ANOVA. *Genotype effect; ^#^ treatment effect. ^#^*P*<0.05; \*\**P*<0.01; ***^,###^*P* < 0.001; ****^,####^*P* < 0.0001.Data are expressed as mean ± S.E.M.

### The characterized CD-associated cardiac reappears after ALO withdrawal

At 6-month-old mice, after two month of ALO withdrawal, we analyzed SBP. All CD (untreated and previously treated) mice shown a significant increased SBP respect to the WT (genotype effect, F_1, 68_ = 111.2; *P*<0.0001) without significant effect of previous treatment (treatment effect F_1, 68_ = 0.2608; *P*=0.6112; Figure 7A).

**FIGURE 7:**
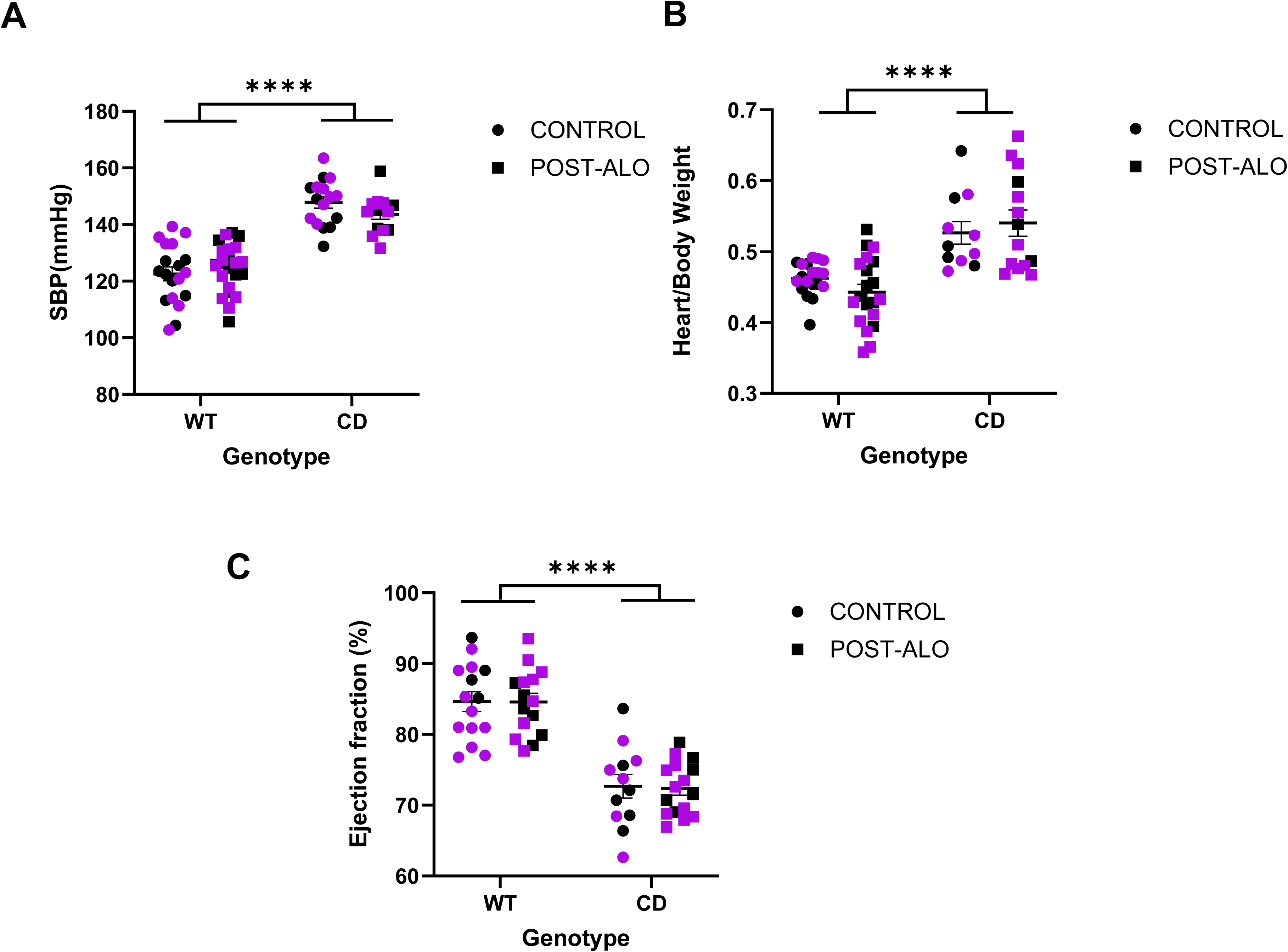
Reappearance of cardiac phenotype in CD mice after ALO treatment withdrawal. **(A)** SBP, **(B)** Heart/Body weigth ratio and **(C)** Ejection Fraction were analyzed from WT and CD 6-month-old mice after 60 days of ALO withdrawal. For all the parameters, we observed a significant genotype effect (*P*<0.0001) without treatment effect, *P*=0.6112 for SBP (A), *P*=0.8256 for heart/body weight ratio (B) and *P*=0.8758 for efection fraction(C). Circles, Control; Squares, Post-ALO treatment. Black, males; Purple, females. Statistical significance was calculated by two-way ANOVA. **** *P*<0.0001 (genotype effect). Data are expressed as mean ± S.E.M.

In relationship with hypercardiopathy, previously treated CD mice presented a significant increase in the heart weight/body weight ratio (genotype effect, F (1,60) = 40.91, *P*<0.0001), without treatment effect (F (1,60) = 0.04898, *P*=0.8256; Figure 7B). Additionally, echocardiography analysis revealed that previously treatred CD mice showed again a significant reduction in EF (genotype effect, F (1,54) = 87.25, *P*<0.0001) independently of treatment (treatment effect F (1, 54) = 0.0246, p=0.8758, Figure 7C). Together, results indicate that ALO withdrawal leadto the reapparence of the characteristic cardiac dysfunction in CD mice. All statistical analysis were detailed in Table S8.

## DISCUSSION

WBS is a distinctive multisystem disorder. We now know that cardiovascular disease is present in most individuals with WBS, with the most frequent abnormalities involving vascular stenoses of medium- and large-sized arteries (referred to as elastin arteriopathy). Although significant progress has been made in understanding WBS since its initial description, numerous challenges persist, particularly regarding the development of effective treatment strategies. Despite the belief that controlling systolic blood pressure (SBP) may mitigate the progression of vascular stiffness and reduce the risk of adverse cardiovascular events, no specific antihypertensive agent has yet been established as the first-line treatment in individuals with WBS [2,12,28].

Here we present a mechanistic study of a clinically available drug (ALO) as a therapeutic alternative in WBS. Elevated XOR expression is a key contributor to the enhanced production of ROS, and its overexpression is linked to increased cardiovascular risk and adverse remodelling in this context [9]. ALO primarily acts as an inhibitor of XOR, a key enzyme in the generation of ROS [29]. ALO has been associated with a small but significant reduction in SBP [30,31]. Therefore, we evaluated the effect of ALO treatment at pediatric doses together with a classic antihypertensive drug (LOS) on the cardiovascular phenotype in WBS model mice [19].

This study produced several main findings: (i) Pediatric dose of LOS and ALO reduce SBP equally eficient;(ii) ALO, as well as LOS, reduces aortic root stenosis in WBS mice; (iii) ALO but not LOS improve the altered cardiac phenotype present in WBS mice; (iv) ALO reduced the WBS-associated increase of XOR expression as well as the subsequent accompanying redox stress-associated reactions such as accumulation of 3-NT; (v) ALO reduced WBS-associated upregulation of NLRP3; (vi) this inhibitory effect was non-permanent since the withdrawal of ALO causes the reappearance of the cardiac phenotype; and (vii) we employed doses that do not affect baseline urate levels [32], but are sufficient to attenuate the oxidative stress–dependent pathology associated with WBS.

Mechanistically, we postulated that pediatric doses of ALO mitigates WBS cardiovascular phenotype acting as an antioxidant both directly scavenging ROS and inhibiting XOR activity, and indirectly downregulating NLRP3 overexpression.

All antihypertensive treatments studied in pediatric WBS patients (calcium channel blockers, renin-angiotensin-aldosterone system inhibitors (angiotensin-converting enzyme inhibitors and angiotensin II type 1 receptor blockers, and beta-blockers) were associated with effective blood pressure management in most cases, although the administration of more than one drug was necessary in 36% of treated patients. [28]. We here demonstrated that both ALO and LOS, at the pediatric doses tested, are equally efficient in reducing increased SBP when administered at 8 weeks of age, when this SBP is already significantly higher in CD animals. Furthermore, although with both treatments, we did not observe changes in the overall diameter of the ascending aorta, they reduced the thickness of the medial wall of these aortas, indicating an improvement in stenosis. This reduction could be attributed to a decrease in cellular hyperplasia of the media.

Surprisingly, seeing the previous results, we observed that treatment with ALO, but not with LOS, improved the cardiac dysfunction and hypertrophy in CD mice. Reversal of cardiac dysfunction and hypertrophy were seen in ALO treated CD mice via improvements in ejection fraction and IVSd (by echocardiografy), and the area of left ventricule and the percentatge of LV muscle (by histology). These improvements are likely due to ALO’s direct inhibition of XOR activity [33]. In line with previous findings [9], we observed a significant increase XOR expression in LV myocardium of CD mice compared to WT mice. ALO treatment normalized XOR expression levels. ALO is a functional inhibitor of XOR and in principle should not affect its protein levels. However, similar reductions have been reported with other functional inhibitors, suggesting that modulation of enzyme activity may, in some cases, influence protein abundance [34]. In agreement with our previous findings [9], associated with the reduction in XOR levels, we also found, a decrease in the levels of nitrosylated proteins (3NT), a characterized biomarker of oxidative stress.

Number of clinical and experimental studies have suggested that XOR activity has pro-inflammatory effects and can mediate cardiovascular and endothelial dysfunction activating NLRP3 inflammasome [25,35,36]. Left ventricular pressure overload triggers NLRP3 inflammasome activation in cardiomyocytes, fibroblasts, and vascular endothelial cells [26].

We have previously demonstrated overexpression of the NLRP3 inflammasome in the cerebral cortex of CD animals [27]. We now demonstrate that this overexpression also occurs in the myocardium and it is normalized after treatment with ALO.

In this context, ALO, an XOR inhibitor widely used in the treatment of hyperuricemia and gout, has emerged as a promising candidate for WBS therapy, and future studies should explore a treatment strategy. This could guide clinical strategies aimed at optimizing treatment regimens, particularly in pediatric populations, where minimizing drug burden and toxicity is critical.

We are aware that our study has some limitations: (i) The timing of treatment initiation (8 weeks) was chosen based on the group’s technical capabilities to monitor the phenotype progression. Although early administration of treatment might induce other adverse effects, a positive preventive effect was reported in a mouse model of Marfan syndrome treated with the same dose of ALO [16]; and (ii) We have only studied the effects of ALO on the cardiovascular system because it is responsible for the life-threatening complication of WBS syndrome. However, since WBS syndrome is a neurodevelopmental disorder, it would also be interesting to study the potential effects of ALO on the neural phenotype, given that it has been shown to cross the blood-brain barrier [37] and is a safe and well-tolerated drug [38].

## Author Contributions

Substantial contributions to the conception/ design of the work: VC-GE

The fine-tuning, acquisition, analysis, drafting, and data interpretation of the work: AAG-IRR-JRC-NA-APD-VC-GE

Revising the work critically for important intellectual content: AAG-IRR-JRC-APD-VC-GE

Final approval of the version to be published: AAG-IRR-JRC-NA-APD-VC-GE

Agreement to be accountable for all aspects of the work in ensuring that questions related to the accuracy or integrity of any part of the work are appropriately investigated and resolved: AAG-IRR-JRC-NA-APD-VC-GE.

For the preparation and writing of this article, the authors declare that no IA application has been used.

## Conflict of interests

The authors declare no conflict of interest.

## Funding

This work was supported by grants from the Ministerio de Ciencia e Innovación [PID2021-124799OB-I00 (AEI/MINEICO/FEDER, UE) to VC and PID2023-146296OB-I00 to GE]; the Association Autour des Williams”/Federation Williams France to VC; and Generalitat de Catalunya [2021 SGR 00029].

## Acknowledgements

We thank Maria Encarnación Palomo for technical assistance.

## SUPPLEMENTAL FIGURE LEGENDS

**FIGURE S1:**
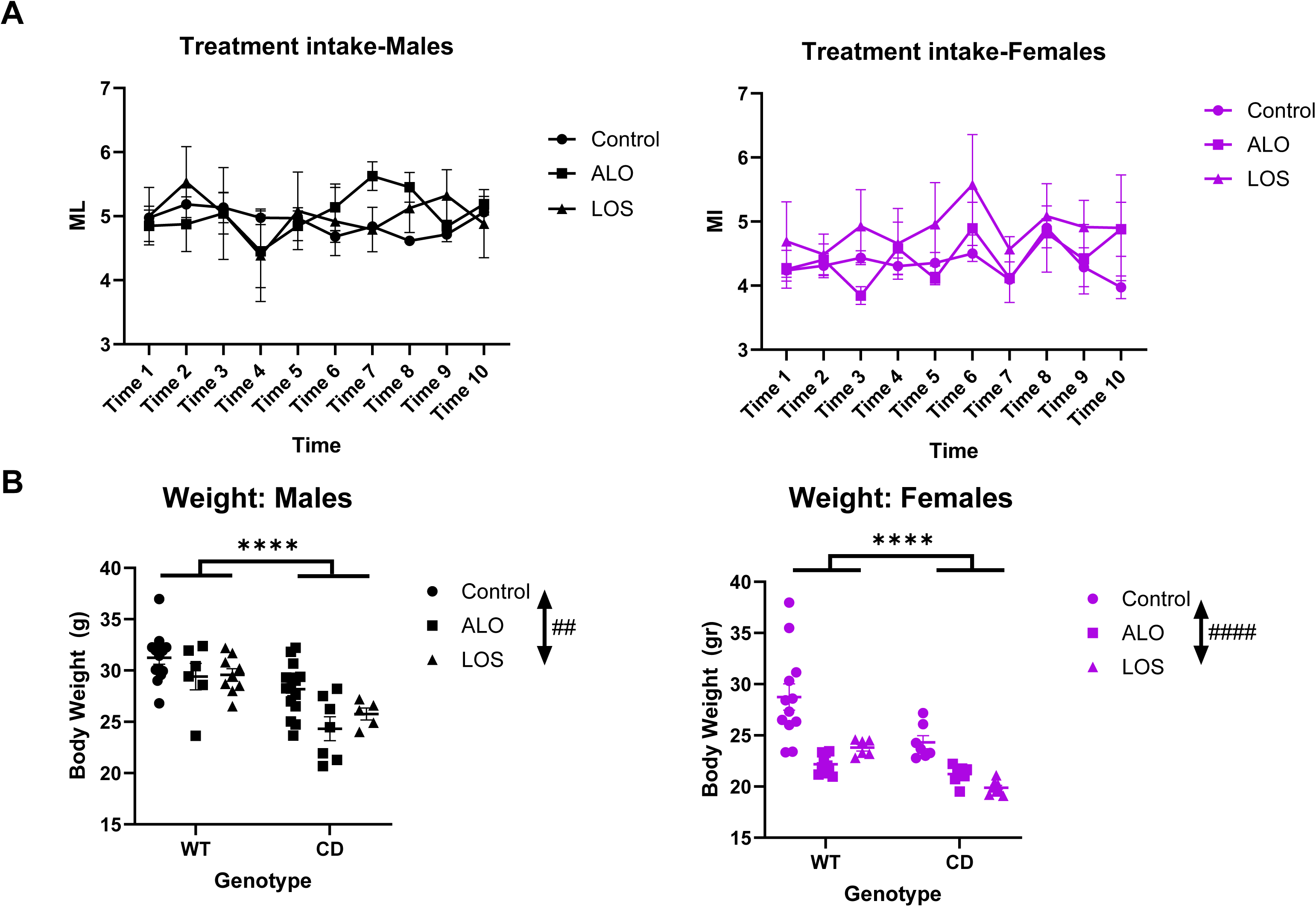
Treatment intake and Body weight. **(A)** Daily consumption (ml/mouse/day) of treatments was equivalent (repeated measures ANOVA, no significant effect of treatment, F_(2,9)_=0.04194, *P*=0.9591 in males (left) and F_(2,9)_=1.208, *P*=0.3431 in females (right)), and did not change over time (two-way ANOVA, no significant interaction between treatment and time, F_(18,81)_=1.652, *P*=0.0666 in males (left) and F_(18,81)_=0.5591, *P*=0.9182 in females(right)). **(B)** Body weights of CD mice were lower in comparison with WT mice without influence of any of the treatments. Two-way ANOVA showed significant effect of genotype (*P*<0.0001) with reduction in the weight of all treated mice effect of treatment (*P*=0.0018 for males and *P*<0,0001 for females). Circles, Control; Squares, ALO treatment. Triangles, LOS treatment. Black, males; Purple, females. *Genotype effect; ^#^ treatment effect. ## *P*<0.01; ****^,####^*P* < 0.0001.Data are expressed as mean ± S.E.M.

**FIGURE S2:**
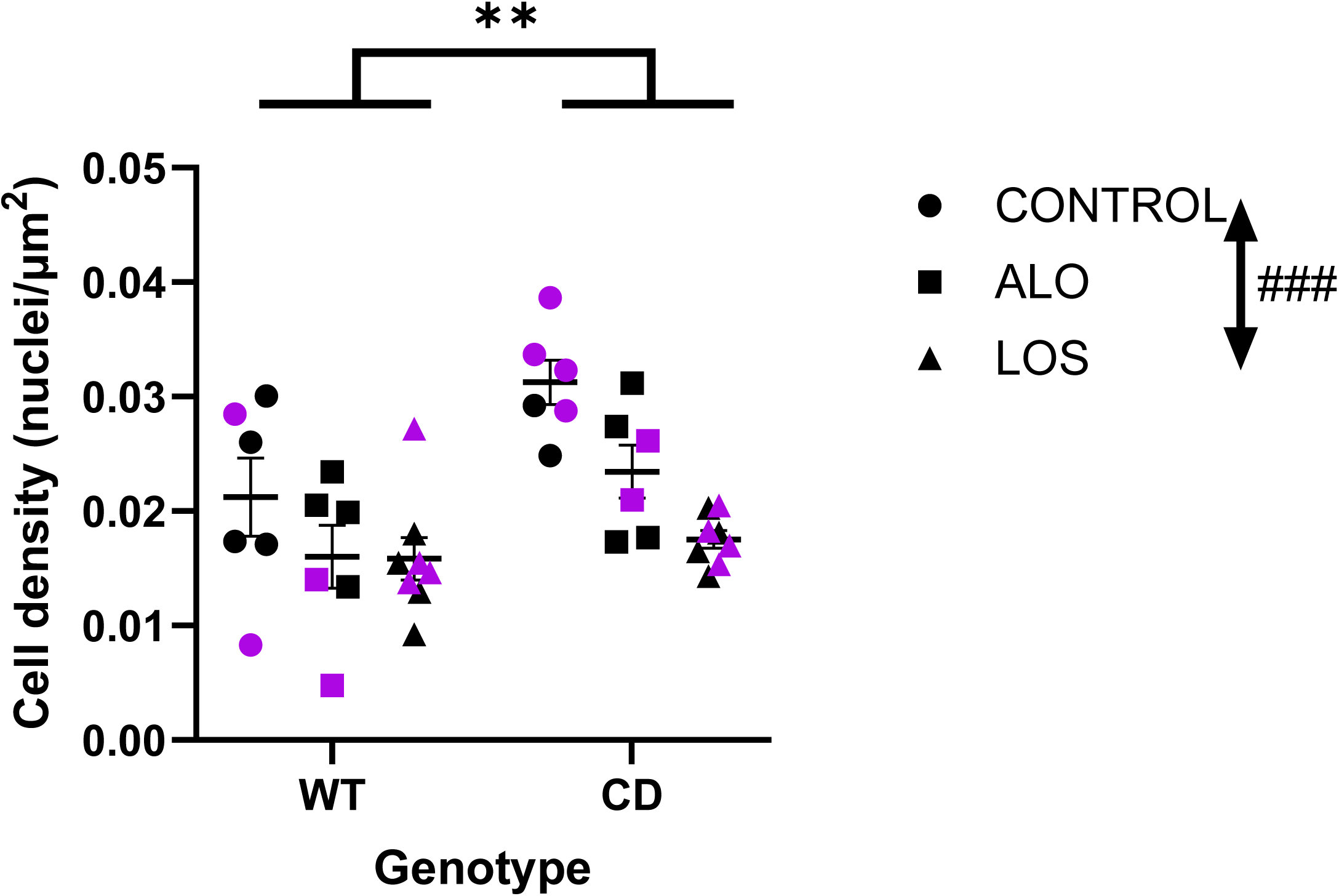
Cell Density. A reduction in cell density (treatment effect, P=0.0004), without genotype interaction, could be observed in all-treated animals (F_(2,34)_=2.031, *P*=0.1468). Statistical significance was calculated by two-way ANOVA. Circles, Control; Squares, ALO treatment; Triangles, LOS treatment. Black, males; Purple, females. *Genotype effect; ^#^ treatment effect. ** *P*<0.01; ^,###^*P* < 0.001.Data are expressed as mean ± S.E.M.

**FIGURE S3:**
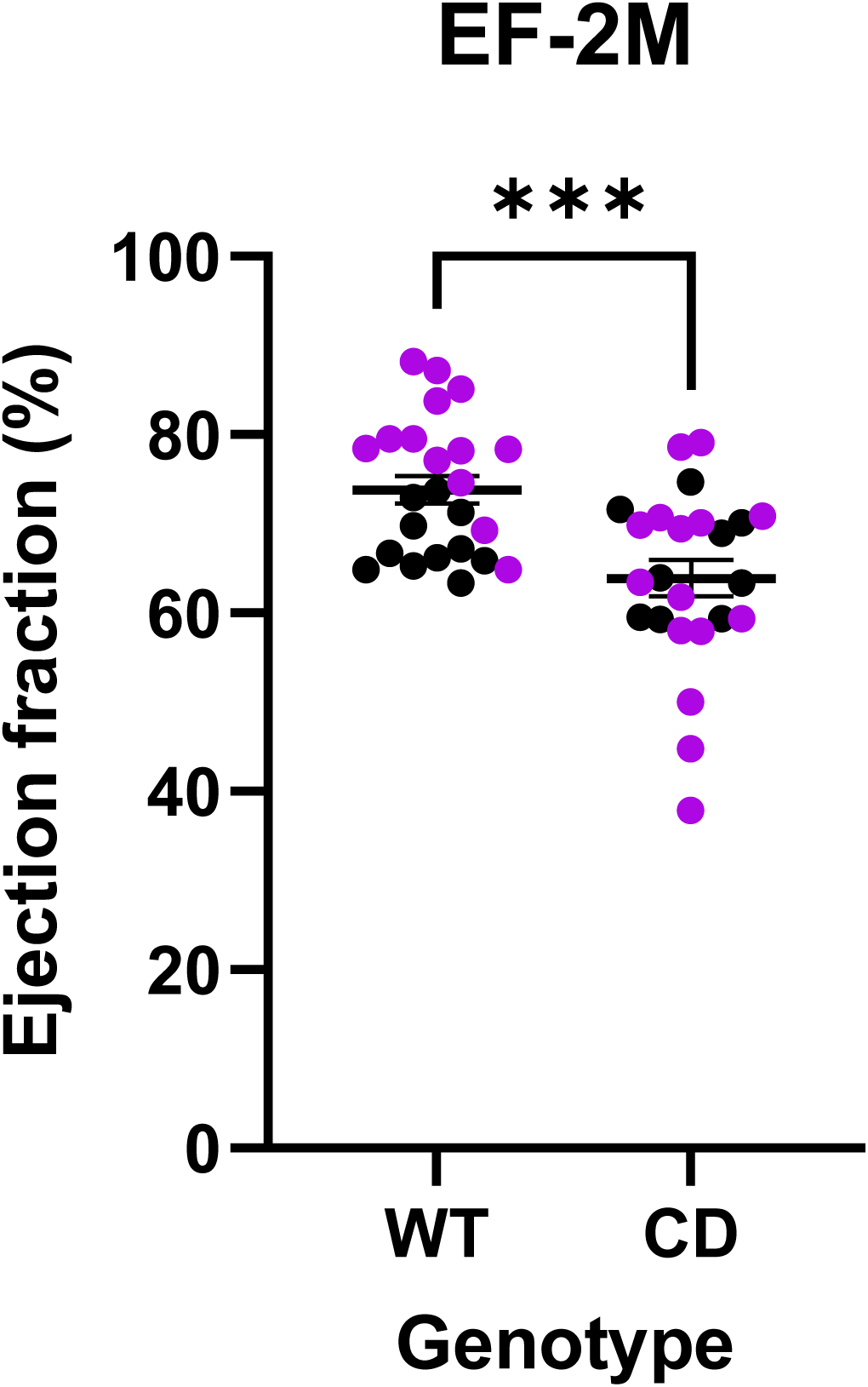
Cardiac dysfunction-associated to CD mice at 2-month-old mice. Ejection Fraction is already reduced in 2-month-old CD mice (P=0.0003). Statistical significance was calculated by Student’s T test. Black, males; Purple, females. *Genotype effect; \*\*\**P* < 0.001

**FIGURE S4:**
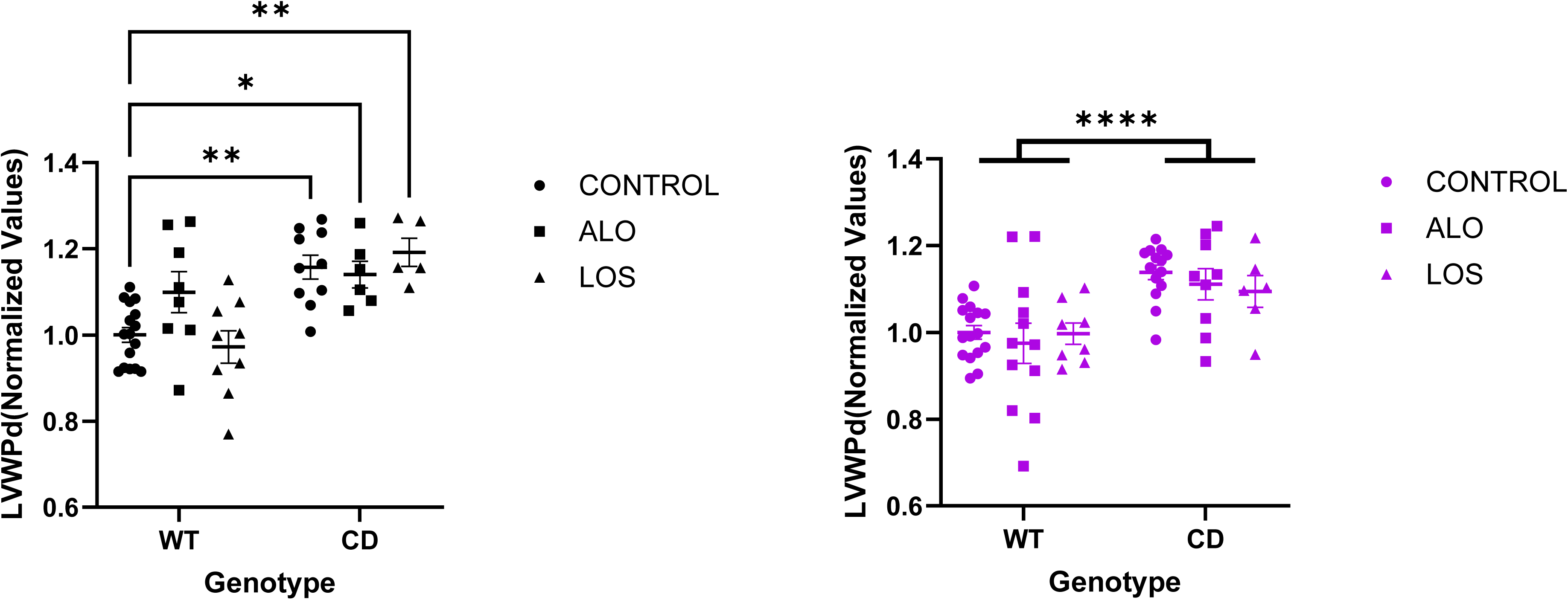
Thickening of the posterior wall of the left ventricle (LVPWd) at 4-month-old mice. Heart hypertrophy in CD animals can be observed in the significant LVPWd thickening (genotype effect, P<0.0001 in males (left) and females (right). LVPWd does not decrease with any of the treatments (treatment effect, P=0.4031 in males and *P*=0.5895 in females). Statistical significance was calculated by two-way ANOVA followed by Tukey’s *post hoc* test in males. Circles, Control; Squares, ALO treatment; Triangles, LOS treatment. Black, males; Purple, females. *Genotype effect; * *P*<0,05; ** *P*<0.01;, \*\*\*\**P* < 0.0001.Data are expressed as mean ± S.E.M.

**FIGURE S5:**
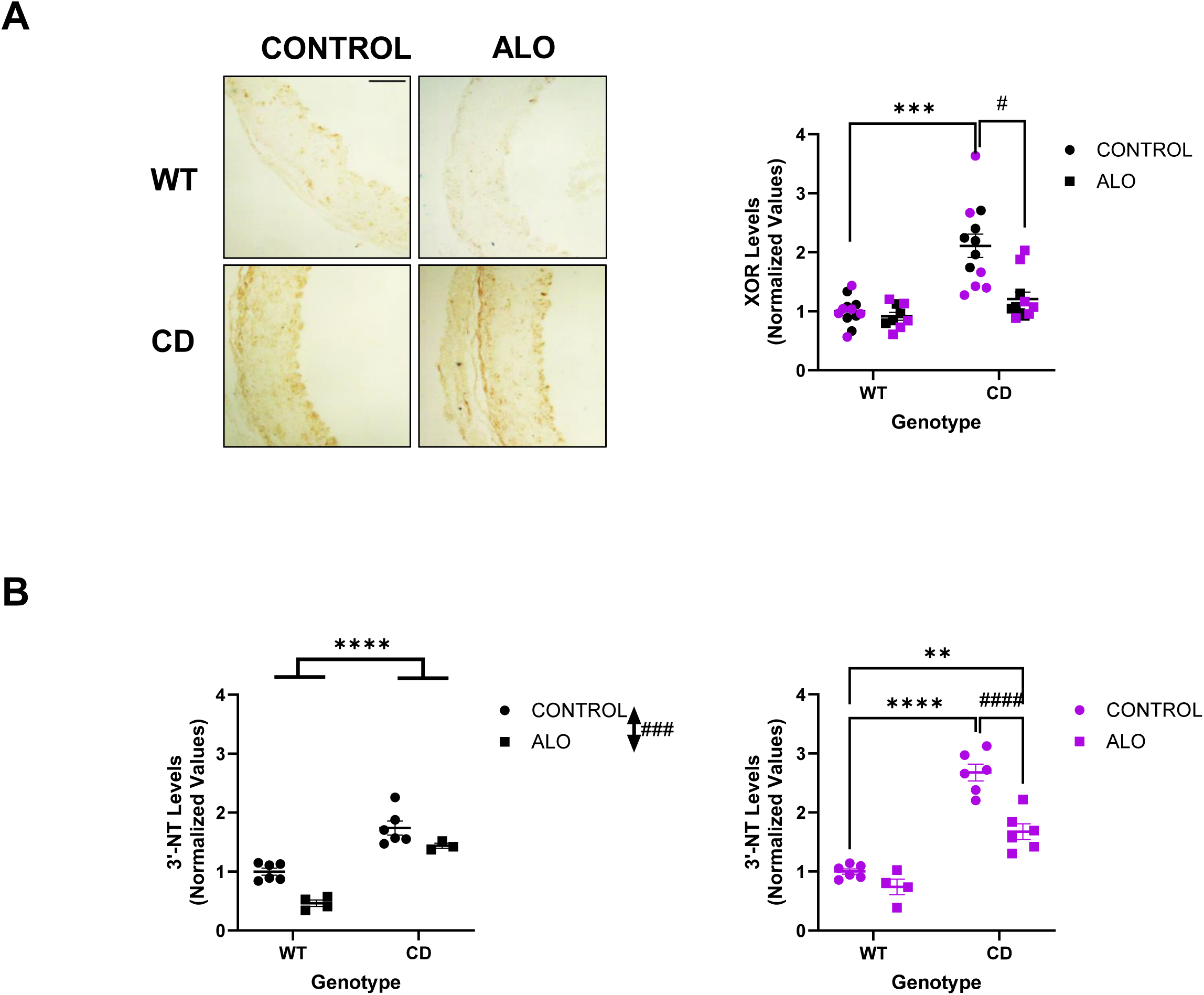
ALO treatment reduces XOR and 3NT levels in ascending aorta of CD mice. **(A)** Representative images of XOR immunohistochemistry detection in ascending aorta (left). Scale bar: 50 µm. A significant increase in XOR protein levels occurs in the ascending aorta of untreated CD mice which were prevented following ALO treatment (*P*>0.9999). **(B).** A significant increase were presented in 3NT levels by ascending aortae of CD mice, both in males and females (genotype effect, *P* < 0.0001. 3NT levels were reduced after ALO treatment (treatment effect, F_(1, 15)_ = 19.88, *P*= 0.0005 in males; F(1, 18) = 26.98, *P*< 0.0001 in females). Values normalized to WT-Control. Circles, Control; Squares, ALO treatment. Black, males; Purple, females. Statistical significance was calculated by Tukey’s *post hoc* test following two-way ANOVA. *Genotype effect; ^#^ treatment effect. ^#^*P*<0.05; \*\**P*<0.01; \*\*\**P* < 0.001; ****^,####^*P* < 0.0001.Data are expressed as mean ± S.E.M.

## SUPPLEMENTAL TABLE

**TABLE S1:** Treatment intake and body weight.

| Treatment intake Males |  |  |  |  |  |  |  |  |  |  |
| --- | --- | --- | --- | --- | --- | --- | --- | --- | --- | --- |
| Shapiro-Wilk test | VEH-T1 | VEH-T2 | VEH-T3 | VEH-T4 | VEH-T5 | VEH-T6 | VEH-T7 | VEH-T8 | VEH-T9 | VEH-T10 |
| W | 0,9774 | 0,9302 | 0,9735 | 0,9443 | 0,8633 | 0,9525 | 0,8655 | 0,9404 | 0,9101 | 0,8834 |
| P value | 0,8869 | 0,5956 | 0,8630 | 0,6808 | 0,2721 | 0,7316 | 0,2804 | 0,6569 | 0,4828 | 0,3534 |
| Shapiro-Wilk test | ALO-T1 | ALO-T2 | ALO-T3 | ALO-T4 | ALO-T5 | ALO-T6 | ALO-T7 | ALO-T8 | ALO-T9 | ALO-T10 |
| W | 0,9529 | 0,9714 | 0,9389 | 0,9343 | 0,7792 | 0,8832 | 0,8945 | 0,8327 | 0,8570 | 0,8634 |
| P value | 0,7341 | 0,8500 | 0,6474 | 0,6198 | 0,0698 | 0,3527 | 0,4043 | 0,1748 | 0,2498 | 0,2725 |
| Shapiro-Wilk test | LOS-T1 | LOS-T2 | LOS-T3 | LOS-T4 | LOS-T5 | LOS-T6 | LOS-T7 | LOS-T8 | LOS-T9 | LOS-T10 |
| W | 0,8955 | 0,9145 | 0,9575 | 0,8235 | 0,8390 | 0,9782 | 0,8737 | 0,9312 | 0,8994 | 0,9245 |
| P value | 0,4090 | 0,5064 | 0,7632 | 0,1514 | 0,1925 | 0,8916 | 0,3126 | 0,6013 | 0,4282 | 0,5626 |
| Treatment intake Females |  |  |  |  |  |  |  |  |  |  |
| Shapiro-Wilk test | VEH-T1 | VEH-T2 | VEH-T3 | VEH-T4 | VEH-T5 | VEH-T6 | VEH-T7 | VEH-T8 | VEH-T9 | VEH-T10 |
| W | 0,9367 | 0,8567 | 0,9306 | 0,9666 | 0,8504 | 0,9498 | 0,8066 | 0,9660 | 0,7674 | 0,9008 |
| P value | 0,6342 | 0,2485 | 0,5981 | 0,8205 | 0,2275 | 0,7147 | 0,1145 | 0,8169 | 0,0555 | 0,4353 |
| Shapiro-Wilk test | ALO-T1 | ALO-T2 | ALO-T3 | ALO-T4 | ALO-T5 | ALO-T6 | ALO-T7 | ALO-T8 | ALO-T9 | ALO-T10 |
| W | 0,9858 | 0,8570 | 0,8621 | 0,8884 | 0,8953 | 0,9058 | 0,9196 | 0,9163 | 0,7805 | 0,9593 |
| P value | 0,9348 | 0,2498 | 0,2678 | 0,3757 | 0,4083 | 0,4603 | 0,5349 | 0,5164 | 0,0715 | 0,7744 |
| Shapiro-Wilk test | LOS-T1 | LOS-T2 | LOS-T3 | LOS-T4 | LOS-T5 | LOS-T6 | LOS-T7 | LOS-T8 | LOS-T9 | LOS-T10 |
| W | 0,8203 | 0,8626 | 0,9215 | 0,8385 | 0,9377 | 0,9299 | 0,9055 | 0,9220 | 0,9145 | 0,8881 |
| P value | 0,1438 | 0,2696 | 0,5456 | 0,1910 | 0,6402 | 0,5939 | 0,4588 | 0,5480 | 0,5066 | 0,3745 |
| Three-way RM ANOVA |  |  | F (DFn, DFd) |  |  |  | P value |  |  |  |
| Time |  |  | F (9, 108) = 1,316 |  |  |  | P=0,2372 |  |  |  |
| SEX |  |  | F (1, 12) = 15,11 |  |  |  | P=0,0022 |  |  |  |
| Treatment |  |  | F (1, 12) = 0,2862 |  |  |  | P=0,6024 |  |  |  |
| Treatment intake Males |  |  |  |  |  |  |  |  |  |  |
| Two-way RM ANOVA |  |  | F (DFn, DFd) |  |  |  | P value |  |  |  |
| Time x Treatment |  |  | F (18, 81) = 1,652 |  |  |  | P=0,0666 |  |  |  |
| Time |  |  | F (1,708, 15,37) = 1,444 |  |  |  | P=0,2636 |  |  |  |
| Treatment |  |  | F (2, 9) = 0,04194 |  |  |  | P=0,9591 |  |  |  |

| Treatment intake Females |  |  |  |  |  |  |
| --- | --- | --- | --- | --- | --- | --- |
| Two-way RM ANOVA |  | F (DFn, DFd) |  |  | P value |  |
| Time x Treatment |  | F (18, 81) = 0,5591 |  |  | P=0,9182 |  |
| Time |  | F (2,907, 26,16) = 1,544 |  |  | P=0,2273 |  |
| Treatment |  | F (2, 9) = 1,208 |  |  | P=0,3431 |  |
| Body Weight Males |  |  |  |  |  |  |
| Shapiro-Wilk test | VEH- WT | ALO-WT | LOS-WT | VEH- CD | ALO-CD | LOS-CD |
| W | 0,9531 | 0,8794 | 0,9779 | 0,9689 | 0,9036 | 0,9352 |
| P value | 0,5753 | 0,2665 | 0,9527 | 0,8413 | 0,3534 | 0,6251 |
| Two-way ANOVA |  | F (DFn, DFd) |  |  | P value |  |
| Interaction |  | F (2, 50) = 0,7606 |  |  | P=0,4727 |  |
| Genotype |  | F (1, 50) = 31,33 |  |  | P<0,0001 |  |
| Treatment |  | F (2, 50) = 7,186 |  |  | P=0,0018 |  |
| Body Weight Females |  |  |  |  |  |  |
| Shapiro-Wilk test | VEH- WT | ALO-WT | LOS-WT | VEH- CD | ALO-CD | LOS-CD |
| W | 0,9154 | 0,8990 | 0,8459 | 0,8600 | 0,9102 | 0,9478 |
| P value | 0,2499 | 0,2134 | 0,1458 | 0,1513 | 0,3555 | 0,6886 |
| Two-way ANOVA |  | F (DFn, DFd) |  |  | P value |  |
| Interaction |  | F (2, 45) = 2,674 |  |  | P=0,0800 |  |
| Genotype |  | F (1, 45) = 20,75 |  |  | P<0,0001 |  |
| Treatment |  | F (2, 45) = 22,97 |  |  | P<0,0001 |  |

**TABLE S2:**
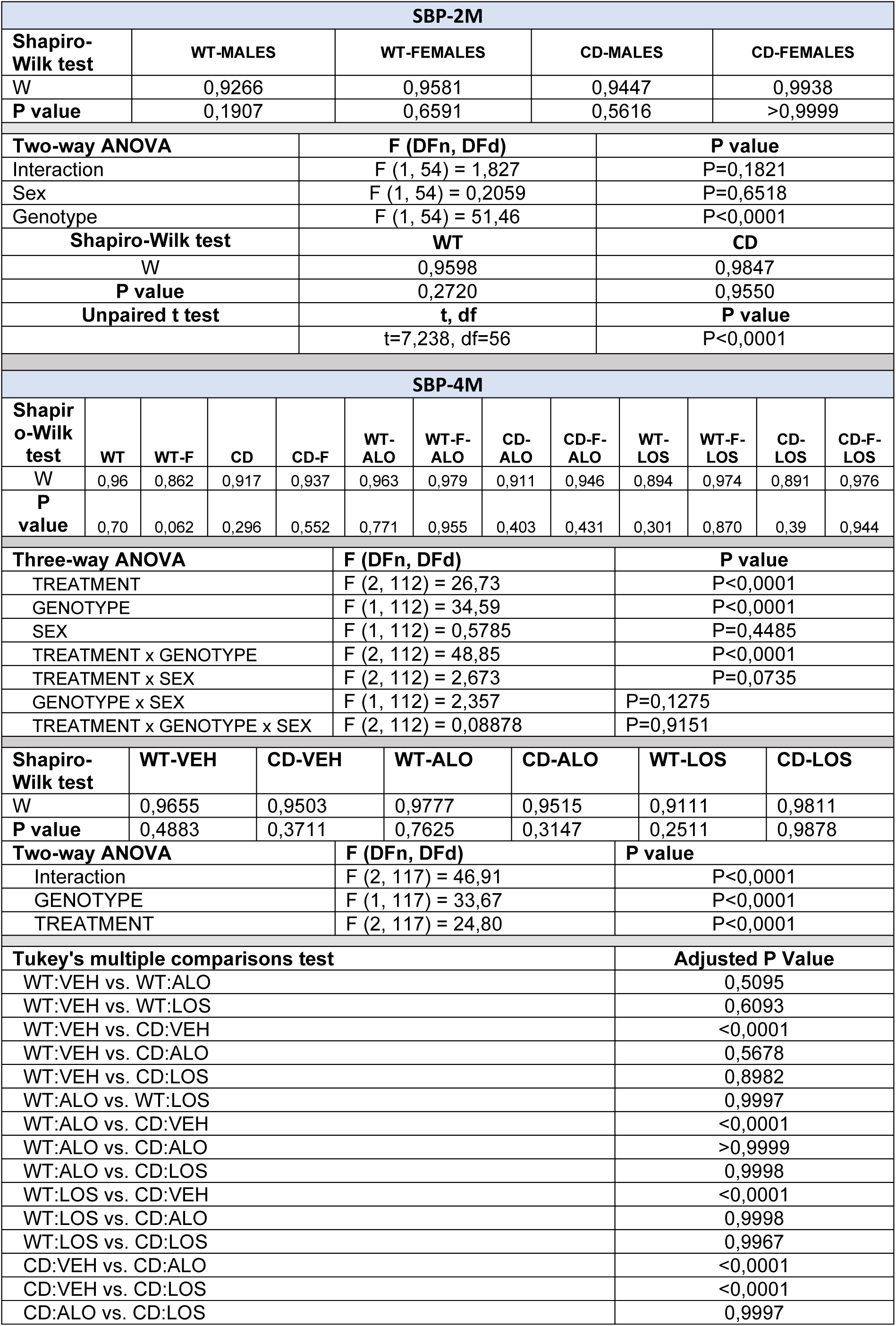
SYS BLOOD PRESSURE.

**TABLE S3:**
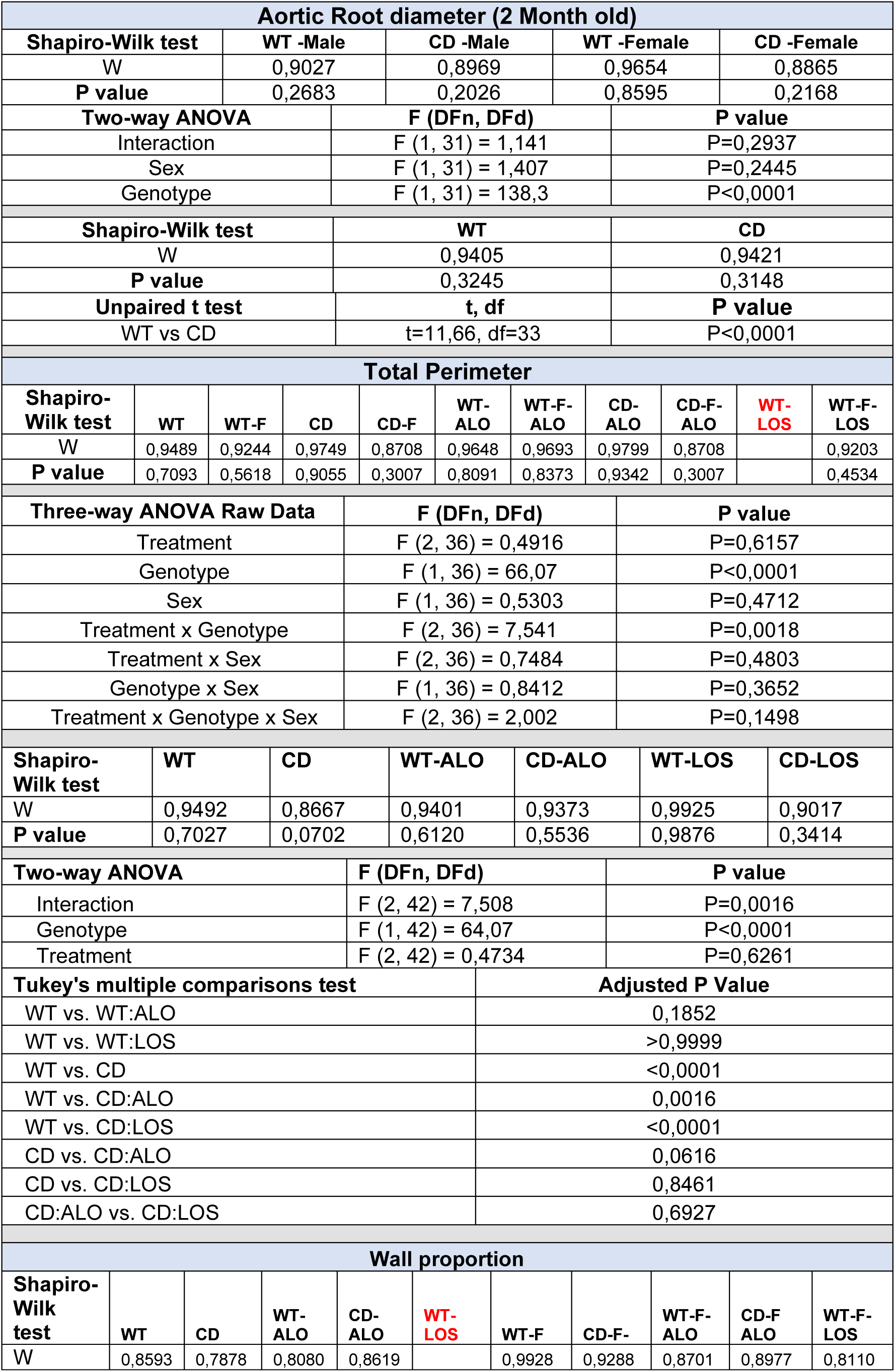
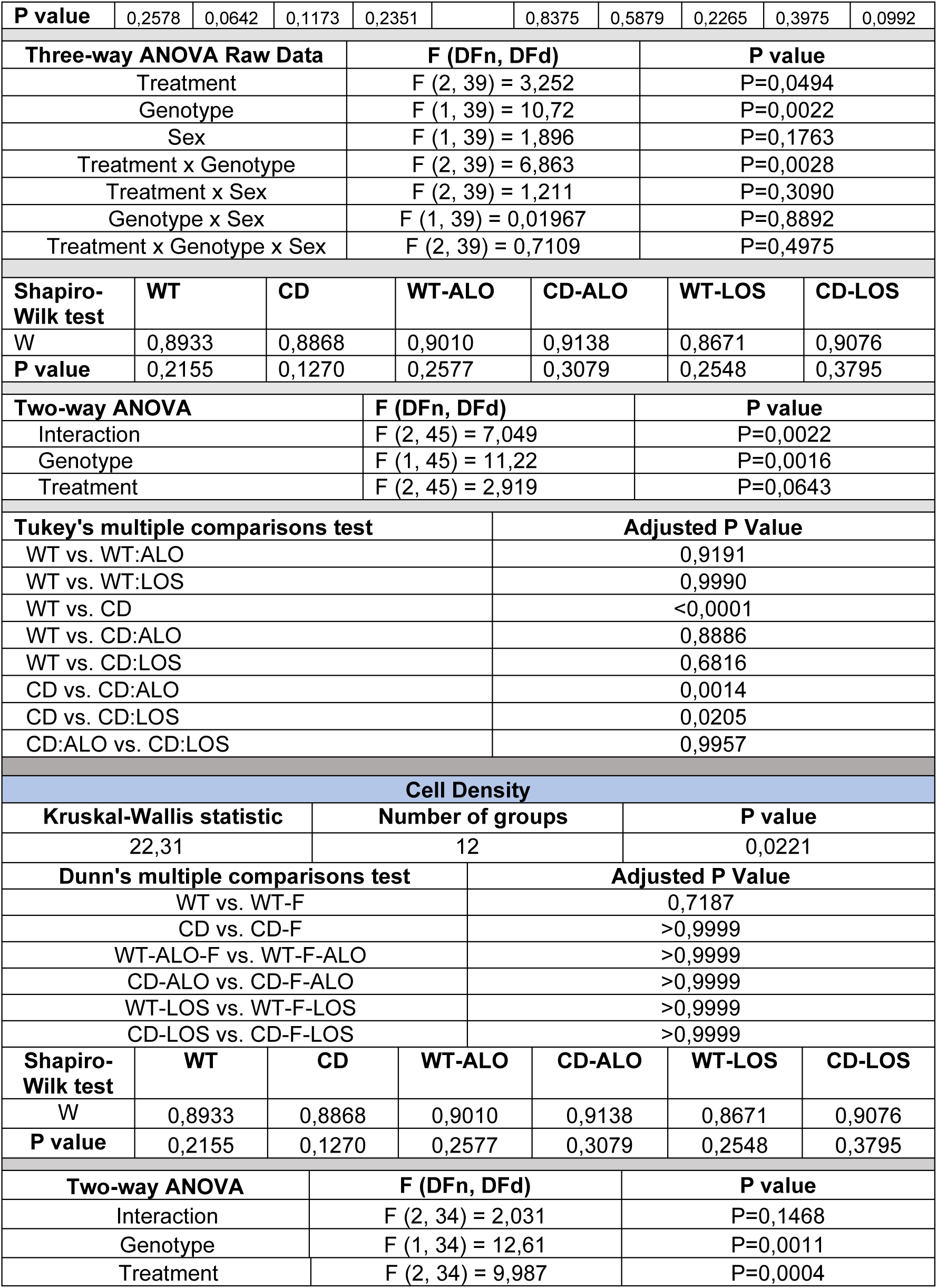
Aortic Root diameter and histomorphometrical parameters in ascending aorta.

| Aortic Root diameter (2 Month old) |  |  |  |  |  |  |  |  |  |  |  |
| --- | --- | --- | --- | --- | --- | --- | --- | --- | --- | --- | --- |
| Shapiro-Wilk test |  | WT -Male |  | CD -Male |  | WT -Female |  | CD -Female |  |  |  |
| W |  | 0,9027 |  | 0,8969 |  | 0,9654 |  | 0,8865 |  |  |  |
| P value |  | 0,2683 |  | 0,2026 |  | 0,8595 |  | 0,2168 |  |  |  |
| Two-way ANOVA |  |  |  | F (DFn, DFd) |  |  |  | P value |  |  |  |
| Interaction |  |  |  | F (1, 31) = 1,141 |  |  |  | P=0,2937 |  |  |  |
| Sex |  |  |  | F (1, 31) = 1,407 |  |  |  | P=0,2445 |  |  |  |
| Genotype |  |  |  | F (1, 31) = 138,3 |  |  |  | P<0,0001 |  |  |  |
| Shapiro-Wilk test |  |  |  | WT |  |  |  | CD |  |  |  |
| W |  |  |  | 0,9405 |  |  |  | 0,9421 |  |  |  |
| P value |  |  |  | 0,3245 |  |  |  | 0,3148 |  |  |  |
| Unpaired t test |  |  |  | t, df |  |  |  | P value |  |  |  |
| WT vs CD |  |  |  | t=11,66, df=33 |  |  |  | P<0,0001 |  |  |  |
| Total Perimeter |  |  |  |  |  |  |  |  |  |  |  |
| Shapiro-Wilk test | WT | WT-F | CD | CD-F | WT-ALO | WT-F-ALO | CD-ALO | CD-F-ALO | WT-LOS | WT-F-LOS |  |
| W | 0,9489 | 0,9244 | 0,9749 | 0,8708 | 0,9648 | 0,9693 | 0,9799 | 0,8708 |  | 0,9203 |  |
| P value | 0,7093 | 0,5618 | 0,9055 | 0,3007 | 0,8091 | 0,8373 | 0,9342 | 0,3007 |  | 0,4534 |  |
| Three-way ANOVA Raw Data |  |  |  | F (DFn, DFd) |  |  |  | P value |  |  |  |
| Treatment |  |  |  | F (2, 36) = 0,4916 |  |  |  | P=0,6157 |  |  |  |
| Genotype |  |  |  | F (1, 36) = 66,07 |  |  |  | P<0,0001 |  |  |  |
| Sex |  |  |  | F (1, 36) = 0,5303 |  |  |  | P=0,4712 |  |  |  |
| Treatment x Genotype |  |  |  | F (2, 36) = 7,541 |  |  |  | P=0,0018 |  |  |  |
| Treatment x Sex |  |  |  | F (2, 36) = 0,7484 |  |  |  | P=0,4803 |  |  |  |
| Genotype x Sex |  |  |  | F (1, 36) = 0,8412 |  |  |  | P=0,3652 |  |  |  |
| Treatment x Genotype x Sex |  |  |  | F (2, 36) = 2,002 |  |  |  | P=0,1498 |  |  |  |
| Shapiro-Wilk test | WT |  | CD |  | WT-ALO |  | CD-ALO |  | WT-LOS |  | CD-LOS |
| W | 0,9492 |  | 0,8667 |  | 0,9401 |  | 0,9373 |  | 0,9925 |  | 0,9017 |
| P value | 0,7027 |  | 0,0702 |  | 0,6120 |  | 0,5536 |  | 0,9876 |  | 0,3414 |
| Two-way ANOVA |  |  |  | F (DFn, DFd) |  |  |  | P value |  |  |  |
| Interaction |  |  |  | F (2, 42) = 7,508 |  |  |  | P=0,0016 |  |  |  |
| Genotype |  |  |  | F (1, 42) = 64,07 |  |  |  | P<0,0001 |  |  |  |
| Treatment |  |  |  | F (2, 42) = 0,4734 |  |  |  | P=0,6261 |  |  |  |
| Tukey's multiple comparisons test |  |  |  |  |  | Adjusted P Value |  |  |  |  |  |
| WT vs. WT:ALO |  |  |  |  |  | 0,1852 |  |  |  |  |  |
| WT vs. WT:LOS |  |  |  |  |  | >0,9999 |  |  |  |  |  |
| WT vs. CD |  |  |  |  |  | <0,0001 |  |  |  |  |  |
| WT vs. CD:ALO |  |  |  |  |  | 0,0016 |  |  |  |  |  |
| WT vs. CD:LOS |  |  |  |  |  | <0,0001 |  |  |  |  |  |
| CD vs. CD:ALO |  |  |  |  |  | 0,0616 |  |  |  |  |  |
| CD vs. CD:LOS |  |  |  |  |  | 0,8461 |  |  |  |  |  |
| CD:ALO vs. CD:LOS |  |  |  |  |  | 0,6927 |  |  |  |  |  |
| Wall proportion |  |  |  |  |  |  |  |  |  |  |  |
| Shapiro-Wilk test | WT | CD | WT-ALO | CD-ALO | WT-LOS | WT-F | CD-F | WT-F-ALO | CD-F-ALO | WT-F-LOS |  |
| W | 0.8593 | 0.7878 | 0.8080 | 0.8619 |  | 0.9928 | 0.9288 | 0.8701 | 0.8977 | 0.8110 |  |

|  |  |  |  |  |  |  |  |  |  |  |
| --- | --- | --- | --- | --- | --- | --- | --- | --- | --- | --- |
| **P value** | 0,2578 | 0,0642 | 0,1173 | 0,2351 |  | 0,8375 | 0,5879 | 0,2265 | 0,3975 | 0,0992 |
| **Three-way ANOVA Raw Data** |  |  | **F (DFn, DFd)** |  |  | **P value** |  |  |  |  |
| Treatment |  |  | F (2, 39) = 3,252 |  |  | P=0,0494 |  |  |  |  |
| Genotype |  |  | F (1, 39) = 10,72 |  |  | P=0,0022 |  |  |  |  |
| Sex |  |  | F (1, 39) = 1,896 |  |  | P=0,1763 |  |  |  |  |
| Treatment x Genotype |  |  | F (2, 39) = 6,863 |  |  | P=0,0028 |  |  |  |  |
| Treatment x Sex |  |  | F (2, 39) = 1,211 |  |  | P=0,3090 |  |  |  |  |
| Genotype x Sex |  |  | F (1, 39) = 0,01967 |  |  | P=0,8892 |  |  |  |  |
| Treatment x Genotype x Sex |  |  | F (2, 39) = 0,7109 |  |  | P=0,4975 |  |  |  |  |
| **Shapiro-Wilk test** | **WT** | **CD** | **WT-ALO** | **CD-ALO** | **WT-LOS** | **CD-LOS** |  |  |  |  |
| W | 0,8933 | 0,8868 | 0,9010 | 0,9138 | 0,8671 | 0,9076 |  |  |  |  |
| **P value** | 0,2155 | 0,1270 | 0,2577 | 0,3079 | 0,2548 | 0,3795 |  |  |  |  |
| **Two-way ANOVA** |  |  | **F (DFn, DFd)** |  |  | **P value** |  |  |  |  |
| Interaction |  |  | F (2, 45) = 7,049 |  |  | P=0,0022 |  |  |  |  |
| Genotype |  |  | F (1, 45) = 11,22 |  |  | P=0,0016 |  |  |  |  |
| Treatment |  |  | F (2, 45) = 2,919 |  |  | P=0,0643 |  |  |  |  |
| **Tukey's multiple comparisons test** |  |  |  | **Adjusted P Value** |  |  |  |  |  |  |
| WT vs. WT:ALO |  |  |  | 0,9191 |  |  |  |  |  |  |
| WT vs. WT:LOS |  |  |  | 0,9990 |  |  |  |  |  |  |
| WT vs. CD |  |  |  | <0,0001 |  |  |  |  |  |  |
| WT vs. CD:ALO |  |  |  | 0,8886 |  |  |  |  |  |  |
| WT vs. CD:LOS |  |  |  | 0,6816 |  |  |  |  |  |  |
| CD vs. CD:ALO |  |  |  | 0,0014 |  |  |  |  |  |  |
| CD vs. CD:LOS |  |  |  | 0,0205 |  |  |  |  |  |  |
| CD:ALO vs. CD:LOS |  |  |  | 0,9957 |  |  |  |  |  |  |
| **Cell Density** |  |  |  |  |  |  |  |  |  |  |
| **Kruskal-Wallis statistic** |  |  | **Number of groups** |  |  | **P value** |  |  |  |  |
| 22,31 |  |  | 12 |  |  | 0,0221 |  |  |  |  |
| **Dunn's multiple comparisons test** |  |  |  | **Adjusted P Value** |  |  |  |  |  |  |
| WT vs. WT-F |  |  |  | 0,7187 |  |  |  |  |  |  |
| CD vs. CD-F |  |  |  | >0,9999 |  |  |  |  |  |  |
| WT-ALO-F vs. WT-F-ALO |  |  |  | >0,9999 |  |  |  |  |  |  |
| CD-ALO vs. CD-F-ALO |  |  |  | >0,9999 |  |  |  |  |  |  |
| WT-LOS vs. WT-F-LOS |  |  |  | >0,9999 |  |  |  |  |  |  |
| CD-LOS vs. CD-F-LOS |  |  |  | >0,9999 |  |  |  |  |  |  |
| **Shapiro-Wilk test** | **WT** | **CD** | **WT-ALO** | **CD-ALO** | **WT-LOS** | **CD-LOS** |  |  |  |  |
| W | 0,8933 | 0,8868 | 0,9010 | 0,9138 | 0,8671 | 0,9076 |  |  |  |  |
| **P value** | 0,2155 | 0,1270 | 0,2577 | 0,3079 | 0,2548 | 0,3795 |  |  |  |  |
| **Two-way ANOVA** |  |  | **F (DFn, DFd)** |  |  | **P value** |  |  |  |  |
| Interaction |  |  | F (2, 34) = 2,031 |  |  | P=0,1468 |  |  |  |  |
| Genotype |  |  | F (1, 34) = 12,61 |  |  | P=0,0011 |  |  |  |  |
| Treatment |  |  | F (2, 34) = 9,987 |  |  | P=0,0004 |  |  |  |  |

**TABLE S4:**
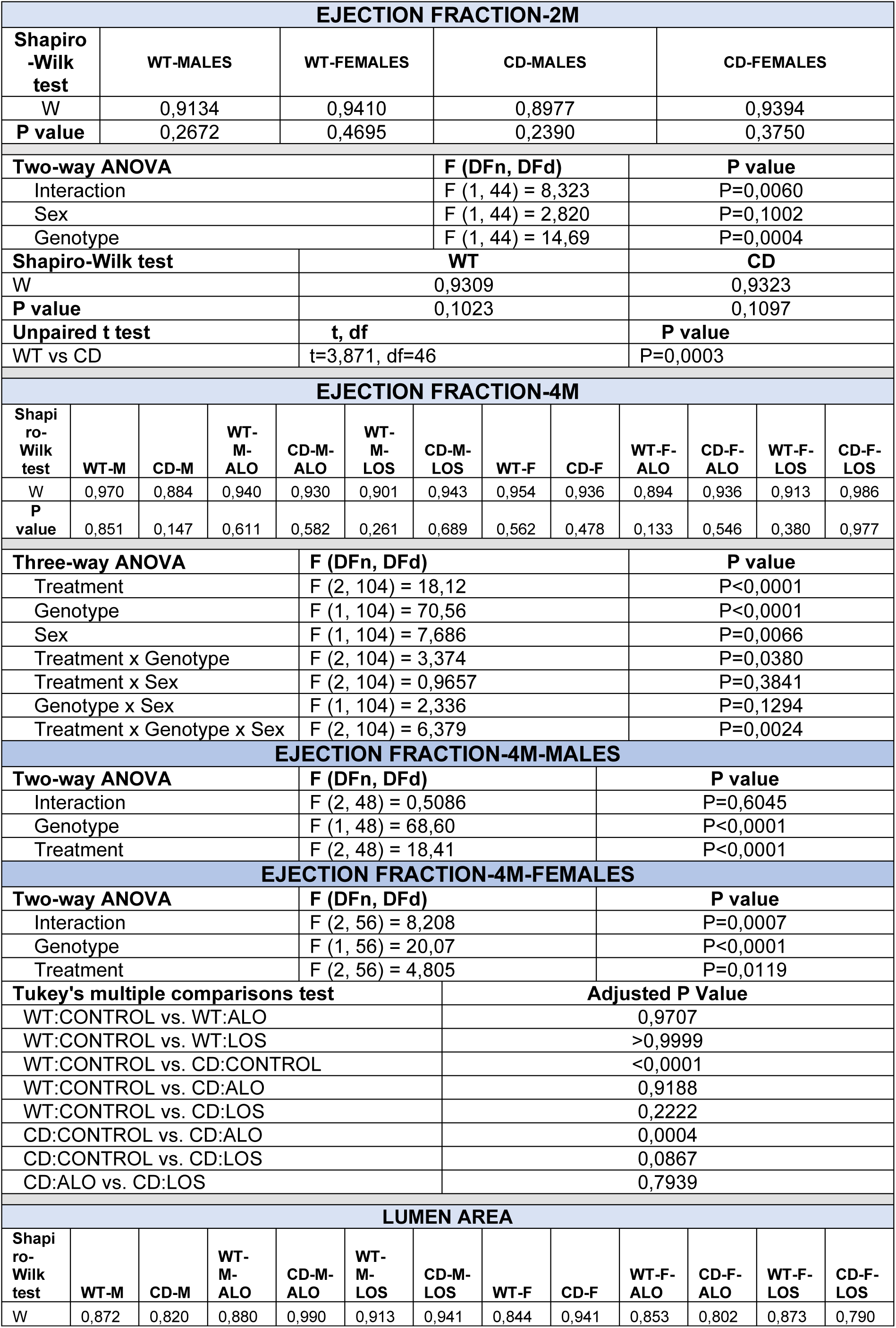
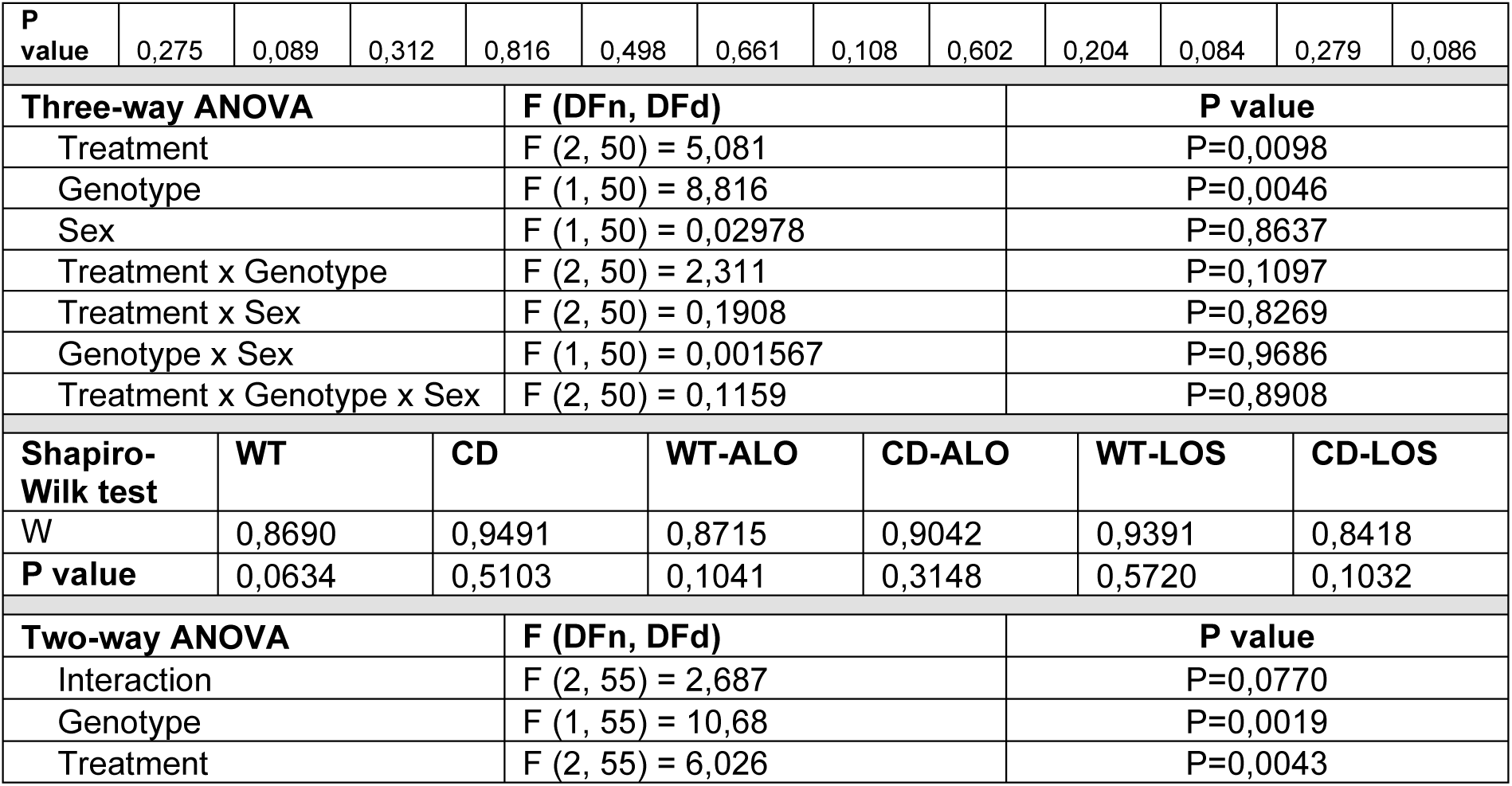
CARDIAC FUNCTION.

| EJECTION FRACTION-2M |  |  |  |  |  |  |  |  |  |  |  |  |
| --- | --- | --- | --- | --- | --- | --- | --- | --- | --- | --- | --- | --- |
| Shapiro-Wilk test | WT-MALES |  |  | WT-FEMALES |  |  | CD-MALES |  |  | CD-FEMALES |  |  |
| W | 0,9134 |  |  | 0,9410 |  |  | 0,8977 |  |  | 0,9394 |  |  |
| P value | 0,2672 |  |  | 0,4695 |  |  | 0,2390 |  |  | 0,3750 |  |  |
| Two-way ANOVA |  |  |  |  |  |  | F (DFn, DFd) |  |  | P value |  |  |
| Interaction |  |  |  |  |  |  | F (1, 44) = 8,323 |  |  | P=0,0060 |  |  |
| Sex |  |  |  |  |  |  | F (1, 44) = 2,820 |  |  | P=0,1002 |  |  |
| Genotype |  |  |  |  |  |  | F (1, 44) = 14,69 |  |  | P=0,0004 |  |  |
| Shapiro-Wilk test |  |  |  | WT |  |  |  |  | CD |  |  |  |
| W |  |  |  | 0,9309 |  |  |  |  | 0,9323 |  |  |  |
| P value |  |  |  | 0,1023 |  |  |  |  | 0,1097 |  |  |  |
| Unpaired t test |  |  |  | t, df |  |  |  |  | P value |  |  |  |
| WT vs CD |  |  |  | t=3,871, df=46 |  |  |  |  | P=0,0003 |  |  |  |
| EJECTION FRACTION-4M |  |  |  |  |  |  |  |  |  |  |  |  |
| Shapiro-Wilk test | WT-M | CD-M | WT-M-ALO | CD-M-ALO | WT-M-LOS | CD-M-LOS | WT-F | CD-F | WT-F-ALO | CD-F-ALO | WT-F-LOS | CD-F-LOS |
| W | 0,970 | 0,884 | 0,940 | 0,930 | 0,901 | 0,943 | 0,954 | 0,936 | 0,894 | 0,936 | 0,913 | 0,986 |
| P value | 0,851 | 0,147 | 0,611 | 0,582 | 0,261 | 0,689 | 0,562 | 0,478 | 0,133 | 0,546 | 0,380 | 0,977 |
| Three-way ANOVA |  |  |  | F (DFn, DFd) |  |  |  |  | P value |  |  |  |
| Treatment |  |  |  | F (2, 104) = 18,12 |  |  |  |  | P<0,0001 |  |  |  |
| Genotype |  |  |  | F (1, 104) = 70,56 |  |  |  |  | P<0,0001 |  |  |  |
| Sex |  |  |  | F (1, 104) = 7,686 |  |  |  |  | P=0,0066 |  |  |  |
| Treatment x Genotype |  |  |  | F (2, 104) = 3,374 |  |  |  |  | P=0,0380 |  |  |  |
| Treatment x Sex |  |  |  | F (2, 104) = 0,9657 |  |  |  |  | P=0,3841 |  |  |  |
| Genotype x Sex |  |  |  | F (1, 104) = 2,336 |  |  |  |  | P=0,1294 |  |  |  |
| Treatment x Genotype x Sex |  |  |  | F (2, 104) = 6,379 |  |  |  |  | P=0,0024 |  |  |  |
| EJECTION FRACTION-4M-MALES |  |  |  |  |  |  |  |  |  |  |  |  |
| Two-way ANOVA |  |  |  | F (DFn, DFd) |  |  |  |  | P value |  |  |  |
| Interaction |  |  |  | F (2, 48) = 0,5086 |  |  |  |  | P=0,6045 |  |  |  |
| Genotype |  |  |  | F (1, 48) = 68,60 |  |  |  |  | P<0,0001 |  |  |  |
| Treatment |  |  |  | F (2, 48) = 18,41 |  |  |  |  | P<0,0001 |  |  |  |
| EJECTION FRACTION-4M-FEMALES |  |  |  |  |  |  |  |  |  |  |  |  |
| Two-way ANOVA |  |  |  | F (DFn, DFd) |  |  |  |  | P value |  |  |  |
| Interaction |  |  |  | F (2, 56) = 8,208 |  |  |  |  | P=0,0007 |  |  |  |
| Genotype |  |  |  | F (1, 56) = 20,07 |  |  |  |  | P<0,0001 |  |  |  |
| Treatment |  |  |  | F (2, 56) = 4,805 |  |  |  |  | P=0,0119 |  |  |  |
| Tukey's multiple comparisons test |  |  |  |  |  |  | Adjusted P Value |  |  |  |  |  |
| WT:CONTROL vs. WT:ALO |  |  |  |  |  |  | 0,9707 |  |  |  |  |  |
| WT:CONTROL vs. WT:LOS |  |  |  |  |  |  | >0,9999 |  |  |  |  |  |
| WT:CONTROL vs. CD:CONTROL |  |  |  |  |  |  | <0,0001 |  |  |  |  |  |
| WT:CONTROL vs. CD:ALO |  |  |  |  |  |  | 0,9188 |  |  |  |  |  |
| WT:CONTROL vs. CD:LOS |  |  |  |  |  |  | 0,2222 |  |  |  |  |  |
| CD:CONTROL vs. CD:ALO |  |  |  |  |  |  | 0,0004 |  |  |  |  |  |
| CD:CONTROL vs. CD:LOS |  |  |  |  |  |  | 0,0867 |  |  |  |  |  |
| CD:ALO vs. CD:LOS |  |  |  |  |  |  | 0,7939 |  |  |  |  |  |
| LUMEN AREA |  |  |  |  |  |  |  |  |  |  |  |  |
| Shapiro-Wilk test | WT-M | CD-M | WT-M-ALO | CD-M-ALO | WT-M-LOS | CD-M-LOS | WT-F | CD-F | WT-F-ALO | CD-F-ALO | WT-F-LOS | CD-F-LOS |
| W | 0,872 | 0,820 | 0,880 | 0,990 | 0,913 | 0,941 | 0,844 | 0,941 | 0,853 | 0,802 | 0,873 | 0,790 |
| **P value** | 0,275 | 0,089 | 0,312 | 0,816 | 0,498 | 0,661 | 0,108 | 0,602 | 0,204 | 0,084 | 0,279 | 0,086 |
| **Three-way ANOVA** |  |  |  | **F (DFn, DFd)** |  |  |  | **P value** |  |  |  |  |
| Treatment |  |  |  | F (2, 50) = 5,081 |  |  |  | P=0,0098 |  |  |  |  |
| Genotype |  |  |  | F (1, 50) = 8,816 |  |  |  | P=0,0046 |  |  |  |  |
| Sex |  |  |  | F (1, 50) = 0,02978 |  |  |  | P=0,8637 |  |  |  |  |
| Treatment x Genotype |  |  |  | F (2, 50) = 2,311 |  |  |  | P=0,1097 |  |  |  |  |
| Treatment x Sex |  |  |  | F (2, 50) = 0,1908 |  |  |  | P=0,8269 |  |  |  |  |
| Genotype x Sex |  |  |  | F (1, 50) = 0,001567 |  |  |  | P=0,9686 |  |  |  |  |
| Treatment x Genotype x Sex |  |  |  | F (2, 50) = 0,1159 |  |  |  | P=0,8908 |  |  |  |  |
| **Shapiro-Wilk test** | **WT** | **CD** | **WT-ALO** |  | **CD-ALO** |  | **WT-LOS** |  | **CD-LOS** |  |  |  |
| W | 0,8690 | 0,9491 | 0,8715 |  | 0,9042 |  | 0,9391 |  | 0,8418 |  |  |  |
| **P value** | 0,0634 | 0,5103 | 0,1041 |  | 0,3148 |  | 0,5720 |  | 0,1032 |  |  |  |
| **Two-way ANOVA** |  |  |  | **F (DFn, DFd)** |  |  |  | **P value** |  |  |  |  |
| Interaction |  |  |  | F (2, 55) = 2,687 |  |  |  | P=0,0770 |  |  |  |  |
| Genotype |  |  |  | F (1, 55) = 10,68 |  |  |  | P=0,0019 |  |  |  |  |
| Treatment |  |  |  | F (2, 55) = 6,026 |  |  |  | P=0,0043 |  |  |  |  |

**TABLE S5:**
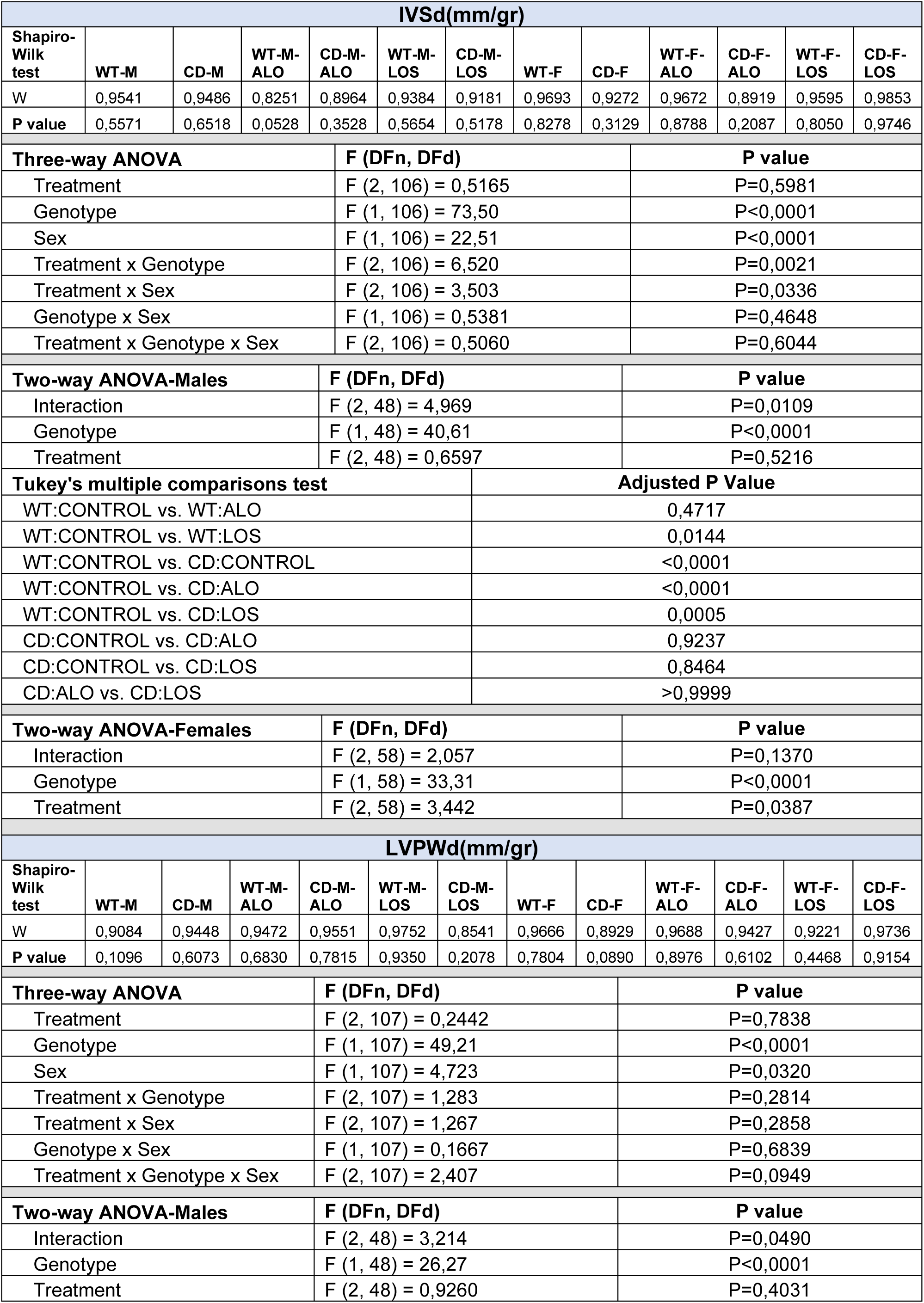
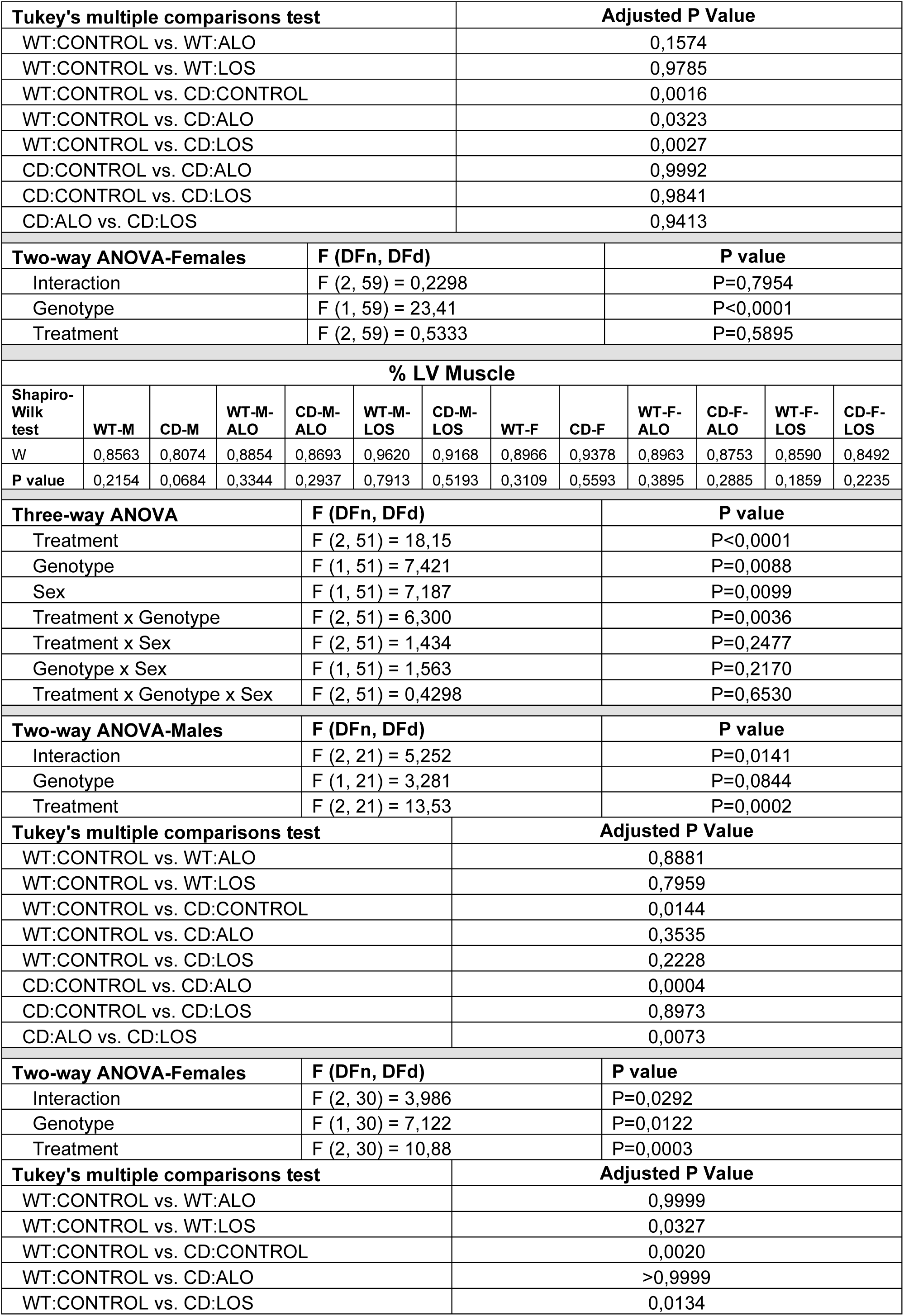
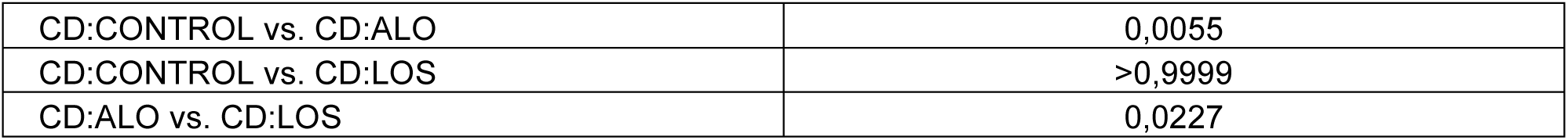
CARDIAC HYPERTROPHY.

| IVSd(mm/gr) |  |  |  |  |  |  |  |  |  |  |  |  |
| --- | --- | --- | --- | --- | --- | --- | --- | --- | --- | --- | --- | --- |
| Shapiro-Wilk test | WT-M | CD-M | WT-M-ALO | CD-M-ALO | WT-M-LOS | CD-M-LOS | WT-F | CD-F | WT-F-ALO | CD-F-ALO | WT-F-LOS | CD-F-LOS |
| W | 0,9541 | 0,9486 | 0,8251 | 0,8964 | 0,9384 | 0,9181 | 0,9693 | 0,9272 | 0,9672 | 0,8919 | 0,9595 | 0,9853 |
| P value | 0,5571 | 0,6518 | 0,0528 | 0,3528 | 0,5654 | 0,5178 | 0,8278 | 0,3129 | 0,8788 | 0,2087 | 0,8050 | 0,9746 |
| Three-way ANOVA |  |  |  | F (DFn, DFd) |  |  |  | P value |  |  |  |  |
| Treatment |  |  |  | F (2, 106) = 0,5165 |  |  |  | P=0,5981 |  |  |  |  |
| Genotype |  |  |  | F (1, 106) = 73,50 |  |  |  | P<0,0001 |  |  |  |  |
| Sex |  |  |  | F (1, 106) = 22,51 |  |  |  | P<0,0001 |  |  |  |  |
| Treatment x Genotype |  |  |  | F (2, 106) = 6,520 |  |  |  | P=0,0021 |  |  |  |  |
| Treatment x Sex |  |  |  | F (2, 106) = 3,503 |  |  |  | P=0,0336 |  |  |  |  |
| Genotype x Sex |  |  |  | F (1, 106) = 0,5381 |  |  |  | P=0,4648 |  |  |  |  |
| Treatment x Genotype x Sex |  |  |  | F (2, 106) = 0,5060 |  |  |  | P=0,6044 |  |  |  |  |
| Two-way ANOVA-Males |  |  |  | F (DFn, DFd) |  |  |  | P value |  |  |  |  |
| Interaction |  |  |  | F (2, 48) = 4,969 |  |  |  | P=0,0109 |  |  |  |  |
| Genotype |  |  |  | F (1, 48) = 40,61 |  |  |  | P<0,0001 |  |  |  |  |
| Treatment |  |  |  | F (2, 48) = 0,6597 |  |  |  | P=0,5216 |  |  |  |  |
| Tukey's multiple comparisons test |  |  |  |  |  | Adjusted P Value |  |  |  |  |  |  |
| WT:CONTROL vs. WT:ALO |  |  |  |  |  | 0,4717 |  |  |  |  |  |  |
| WT:CONTROL vs. WT:LOS |  |  |  |  |  | 0,0144 |  |  |  |  |  |  |
| WT:CONTROL vs. CD:CONTROL |  |  |  |  |  | <0,0001 |  |  |  |  |  |  |
| WT:CONTROL vs. CD:ALO |  |  |  |  |  | <0,0001 |  |  |  |  |  |  |
| WT:CONTROL vs. CD:LOS |  |  |  |  |  | 0,0005 |  |  |  |  |  |  |
| CD:CONTROL vs. CD:ALO |  |  |  |  |  | 0,9237 |  |  |  |  |  |  |
| CD:CONTROL vs. CD:LOS |  |  |  |  |  | 0,8464 |  |  |  |  |  |  |
| CD:ALO vs. CD:LOS |  |  |  |  |  | >0,9999 |  |  |  |  |  |  |
| Two-way ANOVA-Females |  |  |  | F (DFn, DFd) |  |  |  | P value |  |  |  |  |
| Interaction |  |  |  | F (2, 58) = 2,057 |  |  |  | P=0,1370 |  |  |  |  |
| Genotype |  |  |  | F (1, 58) = 33,31 |  |  |  | P<0,0001 |  |  |  |  |
| Treatment |  |  |  | F (2, 58) = 3,442 |  |  |  | P=0,0387 |  |  |  |  |
| LVPWd(mm/gr) |  |  |  |  |  |  |  |  |  |  |  |  |
| Shapiro-Wilk test | WT-M | CD-M | WT-M-ALO | CD-M-ALO | WT-M-LOS | CD-M-LOS | WT-F | CD-F | WT-F-ALO | CD-F-ALO | WT-F-LOS | CD-F-LOS |
| W | 0,9084 | 0,9448 | 0,9472 | 0,9551 | 0,9752 | 0,8541 | 0,9666 | 0,8929 | 0,9688 | 0,9427 | 0,9221 | 0,9736 |
| P value | 0,1096 | 0,6073 | 0,6830 | 0,7815 | 0,9350 | 0,2078 | 0,7804 | 0,0890 | 0,8976 | 0,6102 | 0,4468 | 0,9154 |
| Three-way ANOVA |  |  |  | F (DFn, DFd) |  |  |  | P value |  |  |  |  |
| Treatment |  |  |  | F (2, 107) = 0,2442 |  |  |  | P=0,7838 |  |  |  |  |
| Genotype |  |  |  | F (1, 107) = 49,21 |  |  |  | P<0,0001 |  |  |  |  |
| Sex |  |  |  | F (1, 107) = 4,723 |  |  |  | P=0,0320 |  |  |  |  |
| Treatment x Genotype |  |  |  | F (2, 107) = 1,283 |  |  |  | P=0,2814 |  |  |  |  |
| Treatment x Sex |  |  |  | F (2, 107) = 1,267 |  |  |  | P=0,2858 |  |  |  |  |
| Genotype x Sex |  |  |  | F (1, 107) = 0,1667 |  |  |  | P=0,6839 |  |  |  |  |
| Treatment x Genotype x Sex |  |  |  | F (2, 107) = 2,407 |  |  |  | P=0,0949 |  |  |  |  |
| Two-way ANOVA-Males |  |  |  | F (DFn, DFd) |  |  |  | P value |  |  |  |  |
| Interaction |  |  |  | F (2, 48) = 3,214 |  |  |  | P=0,0490 |  |  |  |  |
| Genotype |  |  |  | F (1, 48) = 26,27 |  |  |  | P<0,0001 |  |  |  |  |
| Treatment |  |  |  | F (2, 48) = 0,9260 |  |  |  | P=0,4031 |  |  |  |  |

| Tukey's multiple comparisons test |  |  |  |  |  |  | Adjusted P Value |  |  |  |  |  |
| --- | --- | --- | --- | --- | --- | --- | --- | --- | --- | --- | --- | --- |
| WT:CONTROL vs. WT:ALO |  |  |  |  |  |  | 0,1574 |  |  |  |  |  |
| WT:CONTROL vs. WT:LOS |  |  |  |  |  |  | 0,9785 |  |  |  |  |  |
| WT:CONTROL vs. CD:CONTROL |  |  |  |  |  |  | 0,0016 |  |  |  |  |  |
| WT:CONTROL vs. CD:ALO |  |  |  |  |  |  | 0,0323 |  |  |  |  |  |
| WT:CONTROL vs. CD:LOS |  |  |  |  |  |  | 0,0027 |  |  |  |  |  |
| CD:CONTROL vs. CD:ALO |  |  |  |  |  |  | 0,9992 |  |  |  |  |  |
| CD:CONTROL vs. CD:LOS |  |  |  |  |  |  | 0,9841 |  |  |  |  |  |
| CD:ALO vs. CD:LOS |  |  |  |  |  |  | 0,9413 |  |  |  |  |  |
| Two-way ANOVA-Females |  |  |  | F (DFn, DFd) |  |  |  | P value |  |  |  |  |
| Interaction |  |  |  | F (2, 59) = 0,2298 |  |  |  | P=0,7954 |  |  |  |  |
| Genotype |  |  |  | F (1, 59) = 23,41 |  |  |  | P<0,0001 |  |  |  |  |
| Treatment |  |  |  | F (2, 59) = 0,5333 |  |  |  | P=0,5895 |  |  |  |  |
| % LV Muscle |  |  |  |  |  |  |  |  |  |  |  |  |
| Shapiro-Wilk test | WT-M | CD-M | WT-M-ALO | CD-M-ALO | WT-M-LOS | CD-M-LOS | WT-F | CD-F | WT-F-ALO | CD-F-ALO | WT-F-LOS | CD-F-LOS |
| W | 0,8563 | 0,8074 | 0,8854 | 0,8693 | 0,9620 | 0,9168 | 0,8966 | 0,9378 | 0,8963 | 0,8753 | 0,8590 | 0,8492 |
| P value | 0,2154 | 0,0684 | 0,3344 | 0,2937 | 0,7913 | 0,5193 | 0,3109 | 0,5593 | 0,3895 | 0,2885 | 0,1859 | 0,2235 |
| Three-way ANOVA |  |  |  | F (DFn, DFd) |  |  |  | P value |  |  |  |  |
| Treatment |  |  |  | F (2, 51) = 18,15 |  |  |  | P<0,0001 |  |  |  |  |
| Genotype |  |  |  | F (1, 51) = 7,421 |  |  |  | P=0,0088 |  |  |  |  |
| Sex |  |  |  | F (1, 51) = 7,187 |  |  |  | P=0,0099 |  |  |  |  |
| Treatment x Genotype |  |  |  | F (2, 51) = 6,300 |  |  |  | P=0,0036 |  |  |  |  |
| Treatment x Sex |  |  |  | F (2, 51) = 1,434 |  |  |  | P=0,2477 |  |  |  |  |
| Genotype x Sex |  |  |  | F (1, 51) = 1,563 |  |  |  | P=0,2170 |  |  |  |  |
| Treatment x Genotype x Sex |  |  |  | F (2, 51) = 0,4298 |  |  |  | P=0,6530 |  |  |  |  |
| Two-way ANOVA-Males |  |  |  | F (DFn, DFd) |  |  |  | P value |  |  |  |  |
| Interaction |  |  |  | F (2, 21) = 5,252 |  |  |  | P=0,0141 |  |  |  |  |
| Genotype |  |  |  | F (1, 21) = 3,281 |  |  |  | P=0,0844 |  |  |  |  |
| Treatment |  |  |  | F (2, 21) = 13,53 |  |  |  | P=0,0002 |  |  |  |  |
| Tukey's multiple comparisons test |  |  |  |  |  |  | Adjusted P Value |  |  |  |  |  |
| WT:CONTROL vs. WT:ALO |  |  |  |  |  |  | 0,8881 |  |  |  |  |  |
| WT:CONTROL vs. WT:LOS |  |  |  |  |  |  | 0,7959 |  |  |  |  |  |
| WT:CONTROL vs. CD:CONTROL |  |  |  |  |  |  | 0,0144 |  |  |  |  |  |
| WT:CONTROL vs. CD:ALO |  |  |  |  |  |  | 0,3535 |  |  |  |  |  |
| WT:CONTROL vs. CD:LOS |  |  |  |  |  |  | 0,2228 |  |  |  |  |  |
| CD:CONTROL vs. CD:ALO |  |  |  |  |  |  | 0,0004 |  |  |  |  |  |
| CD:CONTROL vs. CD:LOS |  |  |  |  |  |  | 0,8973 |  |  |  |  |  |
| CD:ALO vs. CD:LOS |  |  |  |  |  |  | 0,0073 |  |  |  |  |  |
| Two-way ANOVA-Females |  |  |  | F (DFn, DFd) |  |  |  | P value |  |  |  |  |
| Interaction |  |  |  | F (2, 30) = 3,986 |  |  |  | P=0,0292 |  |  |  |  |
| Genotype |  |  |  | F (1, 30) = 7,122 |  |  |  | P=0,0122 |  |  |  |  |
| Treatment |  |  |  | F (2, 30) = 10,88 |  |  |  | P=0,0003 |  |  |  |  |
| Tukey's multiple comparisons test |  |  |  |  |  |  | Adjusted P Value |  |  |  |  |  |
| WT:CONTROL vs. WT:ALO |  |  |  |  |  |  | 0,9999 |  |  |  |  |  |
| WT:CONTROL vs. WT:LOS |  |  |  |  |  |  | 0,0327 |  |  |  |  |  |
| WT:CONTROL vs. CD:CONTROL |  |  |  |  |  |  | 0,0020 |  |  |  |  |  |
| WT:CONTROL vs. CD:ALO |  |  |  |  |  |  | >0,9999 |  |  |  |  |  |
| WT:CONTROL vs. CD:LOS |  |  |  |  |  |  | 0,0134 |  |  |  |  |  |

|  |  |
| --- | --- |
| CD:CONTROL vs. CD:ALO | 0,0055 |
| CD:CONTROL vs. CD:LOS | >0,9999 |
| CD:ALO vs. CD:LOS | 0,0227 |

**TABLE S6:** XOR AND 3-NT LEVELS IN CARDIOVASCULAR TISSUES.

| XOR EXPRESSION IN HEART |  |  |  |  |  |  |  |  |
| --- | --- | --- | --- | --- | --- | --- | --- | --- |
| Shapiro-Wilk test | WT-M | CD-M | WT-M-ALO | CD-M-ALO | WT-F | CD-F | WT-F-ALO | CD-F-ALO |
| W | 0,7718 | 0,8607 | 0,8796 | 0,9979 | 0,9975 | 0,9956 | 0,9311 | 0,9742 |
| P value | 0,0606 | 0,2695 | 0,3371 | 0,9132 | 0,9996 | 0,9994 | 0,6039 | 0,9195 |
| Three-way ANOVA |  |  | F (DFn, DFd) |  |  | P value |  |  |
| Treatment |  |  | F (1, 30) = 26,14 |  |  | P<0,0001 |  |  |
| Genotype |  |  | F (1, 30) = 57,44 |  |  | P<0,0001 |  |  |
| SEX |  |  | F (1, 30) = 0,3343 |  |  | P=0,5674 |  |  |
| Treatment x Genotype |  |  | F (1, 30) = 7,484 |  |  | P=0,0104 |  |  |
| Treatment x SEX |  |  | F (1, 30) = 0,04710 |  |  | P=0,8297 |  |  |
| Genotype x SEX |  |  | F (1, 30) = 0,1898 |  |  | P=0,6662 |  |  |
| Treatment x Genotype x SEX |  |  | F (1, 30) = 0,1438 |  |  | P=0,7072 |  |  |
| Shapiro-Wilk test | WT |  | CD |  | WT-ALO |  | CD-ALO |  |
| W | 0,9602 |  | 0,9816 |  | 0,9351 |  | 0,9482 |  |
| P value | 0,7886 |  | 0,9730 |  | 0,5309 |  | 0,6706 |  |
| Two-way ANOVA |  |  | F (DFn, DFd) |  |  | P value |  |  |
| Interaction |  |  | F (1, 34) = 9,967 |  |  | P=0,0033 |  |  |
| Genotype |  |  | F (1, 34) = 72,92 |  |  | P<0,0001 |  |  |
| Treatment |  |  | F (1, 34) = 33,06 |  |  | P<0,0001 |  |  |
| Tukey's multiple comparisons test |  |  |  | Adjusted P Value |  |  |  |  |
| WT:CONTROL vs. WT:ALO |  |  |  | 0,2758 |  |  |  |  |
| WT:CONTROL vs. CD:CONTROL |  |  |  | <0,0001 |  |  |  |  |
| WT:CONTROL vs. CD:ALO |  |  |  | 0,2181 |  |  |  |  |
| WT:ALO vs. CD:CONTROL |  |  |  | <0,0001 |  |  |  |  |
| WT:ALO vs. CD:ALO |  |  |  | 0,0039 |  |  |  |  |
| CD:CONTROL vs. CD:ALO |  |  |  | <0,0001 |  |  |  |  |
| XOR EXPRESSION IN AORTA |  |  |  |  |  |  |  |  |
| Shapiro-Wilk test | WT-M | CD-M | WT-M-ALO | CD-M-ALO | WT-F | CD-F | WT-F-ALO | CD-F-ALO |
| W | 0,9833 | 0,9899 | 0,9366 | 0,8955 | 0,9002 | 0,8062 | 0,9197 | 0,8260 |
| P value | 0,9667 | 0,9888 | 0,6339 | 0,3854 | 0,3754 | 0,0668 | 0,5280 | 0,0994 |
| Three-way ANOVA |  |  | F (DFn, DFd) |  |  | P value |  |  |
| Treatment |  |  | F (1, 36) = 10,61 |  |  | P=0,0025 |  |  |
| Genotype |  |  | F (1, 36) = 27,67 |  |  | P<0,0001 |  |  |
| SEX |  |  | F (1, 36) = 0,1374 |  |  | P=0,7131 |  |  |
| Treatment x Genotype |  |  | F (1, 36) = 7,112 |  |  | P=0,0114 |  |  |
| Treatment x SEX |  |  | F (1, 36) = 1,163 |  |  | P=0,2880 |  |  |
| Genotype x SEX |  |  | F (1, 36) = 0,2272 |  |  | P=0,6365 |  |  |
| Treatment x Genotype x SEX |  |  | F (1, 36) = 1,403 |  |  | P=0,2440 |  |  |
| Shapiro-Wilk test | WT |  | CD |  | WT-ALO |  | CD-ALO |  |
| W | 0,9555 |  | 0,9337 |  | 0,9424 |  | 0,7554 |  |
| P value | 0,7182 |  | 0,4215 |  | 0,6072 |  | 0,0024 |  |
| Kruskal-Wallis statistic |  |  | Number of groups |  |  | P value |  |  |
| 23,85 |  |  | 4 |  |  | P<0.0001 |  |  |

| Dunn's multiple comparisons test |  |  |  |  | Adjusted P Value |  |  |  |
| --- | --- | --- | --- | --- | --- | --- | --- | --- |
| WT:CONTROL vs. CD:CONTROL |  |  |  |  | 0,0004 |  |  |  |
| WT:CONTROL vs. WT-ALO |  |  |  |  | >0,9999 |  |  |  |
| WT:CONTROL vs. CD-ALO |  |  |  |  | >0,9999 |  |  |  |
| CD:CONTROL vs. WT-ALO |  |  |  |  | <0,0001 |  |  |  |
| CD:CONTROL vs. CD-ALO |  |  |  |  | 0,0192 |  |  |  |
| WT-ALO vs. CD-ALO |  |  |  |  | 0,7918 |  |  |  |
| 3-NT EXPRESSION IN HEART |  |  |  |  |  |  |  |  |
| Shapiro-Wilk test | WT-M | CD-M | WT-M-ALO | CD-M-ALO | WT-F | CD-F | WT-F-ALO | CD-F-ALO |
| W | 0,8593 | 0,9947 | 0,9102 | 0,9826 | 0,8817 | 0,9557 | 0,7993 | 0,8977 |
| P value | 0,2656 | 0,8606 | 0,4186 | 0,7473 | 0,3293 | 0,7521 | 0,1129 | 0,4198 |
| Three-way ANOVA |  |  | F (DFn, DFd) |  |  | P value |  |  |
| Treatment |  |  | F (1, 18) = 23,38 |  |  | P=0,0001 |  |  |
| Genotype |  |  | F (1, 18) = 26,05 |  |  | P<0,0001 |  |  |
| SEX |  |  | F (1, 18) = 2,635 |  |  | P=0,1219 |  |  |
| Treatment x Genotype |  |  | F (1, 18) = 5,921 |  |  | P=0,0256 |  |  |
| Treatment x SEX |  |  | F (1, 18) = 0,4297 |  |  | P=0,5204 |  |  |
| Genotype x SEX |  |  | F (1, 18) = 0,09410 |  |  | P=0,7625 |  |  |
| Treatment x Genotype x SEX |  |  | F (1, 18) = 0,4368 |  |  | P=0,5170 |  |  |
| Shapiro-Wilk test |  | WT |  | CD |  | WT-ALO |  | CD-ALO |
| W |  | 0,9251 |  | 0,8593 |  | 0,9134 |  | 0,8894 |
| P value |  | 0,5429 |  | 0,1493 |  | 0,4590 |  | 0,2716 |
| Two-way ANOVA |  |  | F (DFn, DFd) |  |  | P value |  |  |
| Interaction |  |  | F (1, 22) = 6,084 |  |  | P=0,0219 |  |  |
| Genotype |  |  | F (1, 22) = 28,23 |  |  | P<0,0001 |  |  |
| Treatment |  |  | F (1, 22) = 24,02 |  |  | P<0,0001 |  |  |
| Tukey's multiple comparisons test |  |  |  |  | Adjusted P Value |  |  |  |
| WT:CONTROL vs. WT:ALO |  |  |  |  | 0,3683 |  |  |  |
| WT:CONTROL vs. CD:CONTROL |  |  |  |  | <0,0001 |  |  |  |
| WT:CONTROL vs. CD:ALO |  |  |  |  | 0,9911 |  |  |  |
| WT:ALO vs. CD:CONTROL |  |  |  |  | <0,0001 |  |  |  |
| WT:ALO vs. CD:ALO |  |  |  |  | 0,2137 |  |  |  |
| CD:CONTROL vs. CD:ALO |  |  |  |  | 0,0001 |  |  |  |
| 3-NT EXPRESSION IN AORTA |  |  |  |  |  |  |  |  |
| Shapiro-Wilk test | WT-M | CD-M | WT-M-ALO | CD-M-ALO | WT-F | CD-F | WT-F-ALO | CD-F-ALO |
| W | 0,8050 | 0,8726 | 0,9438 | 0,9676 | 0,9350 | 0,9697 | 0,9645 | 0,9579 |
| P value | 0,0652 | 0,2366 | 0,6775 | 0,6546 | 0,6193 | 0,8902 | 0,8073 | 0,8038 |
| Three-way ANOVA |  |  | F (DFn, DFd) |  |  | P value |  |  |
| Treatment |  |  | F (1, 33) = 43,76 |  |  | P<0,0001 |  |  |
| Genotype |  |  | F (1, 33) = 186,5 |  |  | P<0,0001 |  |  |
| SEX |  |  | F (1, 33) = 21,09 |  |  | P<0,0001 |  |  |
| Treatment x Genotype |  |  | F (1, 33) = 2,567 |  |  | P=0,1186 |  |  |
| Treatment x SEX |  |  | F (1, 33) = 1,787 |  |  | P=0,1904 |  |  |
| Genotype x SEX |  |  | F (1, 33) = 8,020 |  |  | P=0,0078 |  |  |
| Treatment x Genotype x SEX |  |  | F (1, 33) = 9,460 |  |  | P=0,0042 |  |  |
| 3-NT EXPRESSION IN AORTA-MALES |  |  |  |  |  |  |  |  |

| Two-way ANOVA | F (DFn, DFd) | P value |
| --- | --- | --- |
| Interaction | F (1, 15) = 1,549 | P=0,2324 |
| Genotype | F (1, 15) = 83,56 | P<0,0001 |
| Treatment | F (1, 15) = 19,88 | P=0,0005 |
| 3-NT EXPRESSION IN AORTA-FEMALES |  |  |
| Two-way ANOVA | F (DFn, DFd) | P value |
| Interaction | F (1, 18) = 9,335 | P=0,0068 |
| Genotype | F (1, 18) = 116,0 | P<0,0001 |
| Treatment | F (1, 18) = 26,98 | P<0,0001 |
| Tukey's multiple comparisons test |  | Adjusted P Value |
| WT:CONTROL vs. WT:ALO |  | 0,4952 |
| WT:CONTROL vs. CD:CONTROL |  | <0,0001 |
| WT:CONTROL vs. CD:ALO |  | 0,0029 |
| WT:ALO vs. CD:CONTROL |  | <0,0001 |
| WT:ALO vs. CD:ALO |  | 0,0003 |
| CD:CONTROL vs. CD:ALO |  | <0,0001 |

**TABLE S7:** NRLP3 LEVELS IN MYOCARDIUM.

| NRLP3 LEVELS IN MYOCARDIUM |  |  |  |  |  |  |  |  |
| --- | --- | --- | --- | --- | --- | --- | --- | --- |
| Shapiro-Wilk test | WT-M | CD-M | WT-M-ALO | CD-M-ALO | WT-F | CD-F | WT-F-ALO | CD-F-ALO |
| W | 0,9895 | 0,8342 | 0,8628 | N too small | 0,9909 | 0,9308 | 0,9422 | 0,8906 |
| P value | 0,9550 | 0,1990 | 0,2754 |  | 0,9912 | 0,5576 | 0,6817 | 0,3212 |
| Kruskal-Wallis statistic |  |  | Number of groups |  |  | P value |  |  |
| 25,59 |  |  | 8 |  |  | 0,0006 |  |  |
| Dunn's múltiple comparisons test |  |  |  | Adjusted P Value |  |  |  |  |
| WT.Male vs WT. Female |  |  |  | >0,9999 |  |  |  |  |
| CD.Male vs CD. Female |  |  |  | >0,9999 |  |  |  |  |
| WT.ALO.Male vs WT.ALO. Female |  |  |  | >0,9999 |  |  |  |  |
| CD.ALO.Male vs CD.ALO. Female |  |  |  | >0,9999 |  |  |  |  |
| Shapiro-Wilk test | WT |  | CD |  | WT-ALO |  | CD-ALO |  |
| W | 0,9618 |  | 0,8938 |  | 0,9635 |  | 0,8845 |  |
| P value | 0,8059 |  | 0,1872 |  | 0,8423 |  | 0,2080 |  |
| Two-way ANOVA |  |  | F (DFn, DFd) |  |  | P value |  |  |
| Interaction |  |  | F (1, 32) = 6,668 |  |  | P=0,0146 |  |  |
| Genotype |  |  | F (1, 32) = 32,47 |  |  | P<0,0001 |  |  |
| Treatment |  |  | F (1, 32) = 21,69 |  |  | P<0,0001 |  |  |
| Tukey's multiple comparisons test |  |  |  | Adjusted P Value |  |  |  |  |
| WT:CONTROL vs. WT:ALO |  |  |  | 0,4684 |  |  |  |  |
| WT:CONTROL vs. CD:CONTROL |  |  |  | <0,0001 |  |  |  |  |
| WT:CONTROL vs. CD:ALO |  |  |  | 0,8817 |  |  |  |  |
| WT:ALO vs. CD:CONTROL |  |  |  | <0,0001 |  |  |  |  |
| WT:ALO vs. CD:ALO |  |  |  | 0,1779 |  |  |  |  |
| CD:CONTROL vs. CD:ALO |  |  |  | <0,0001 |  |  |  |  |

**TABLE S8:**
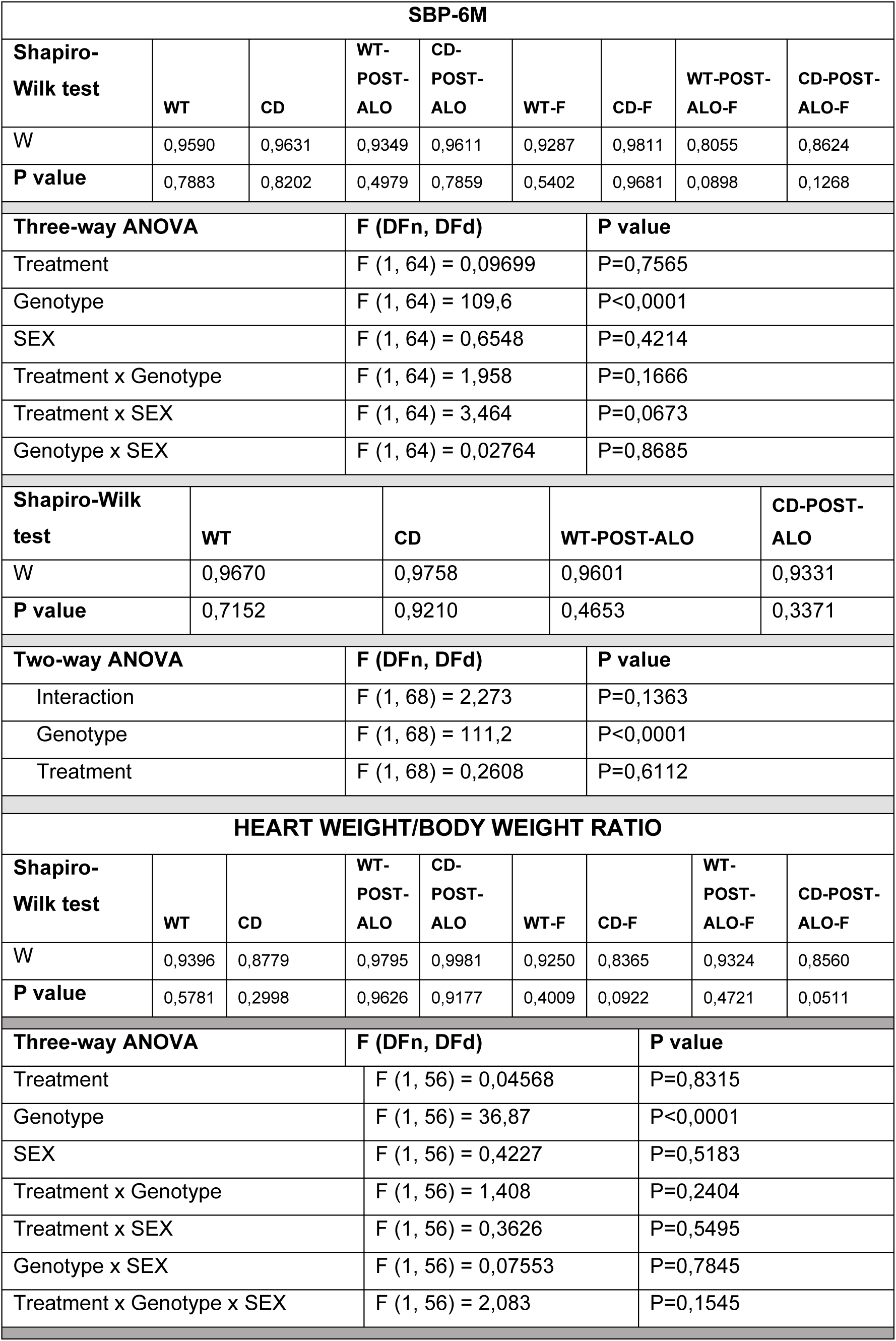
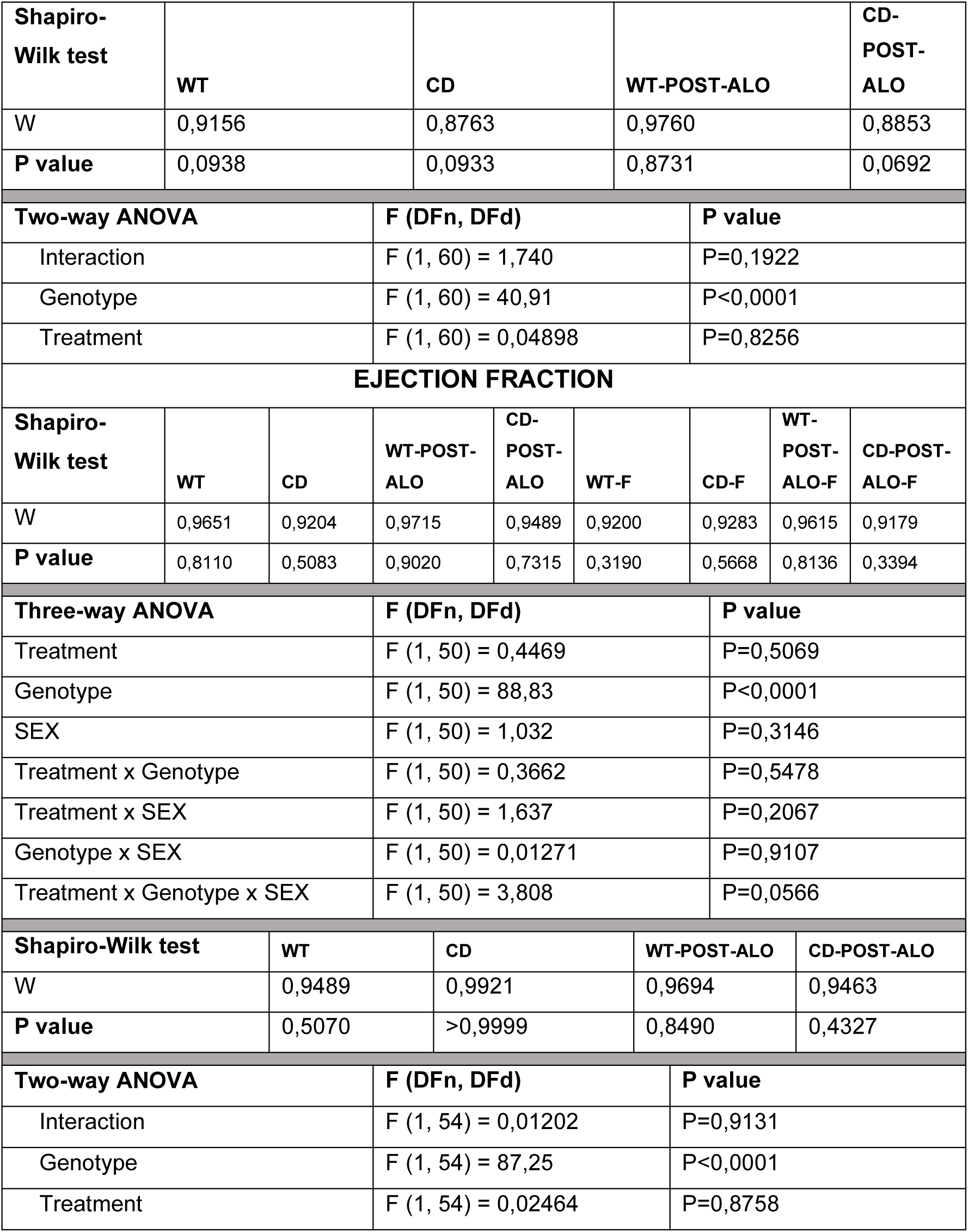
CARDIAVASCULAR PARAMETERS AFTER ALO TREATMENT WITHDRAWAL.

**TABLE S7:**
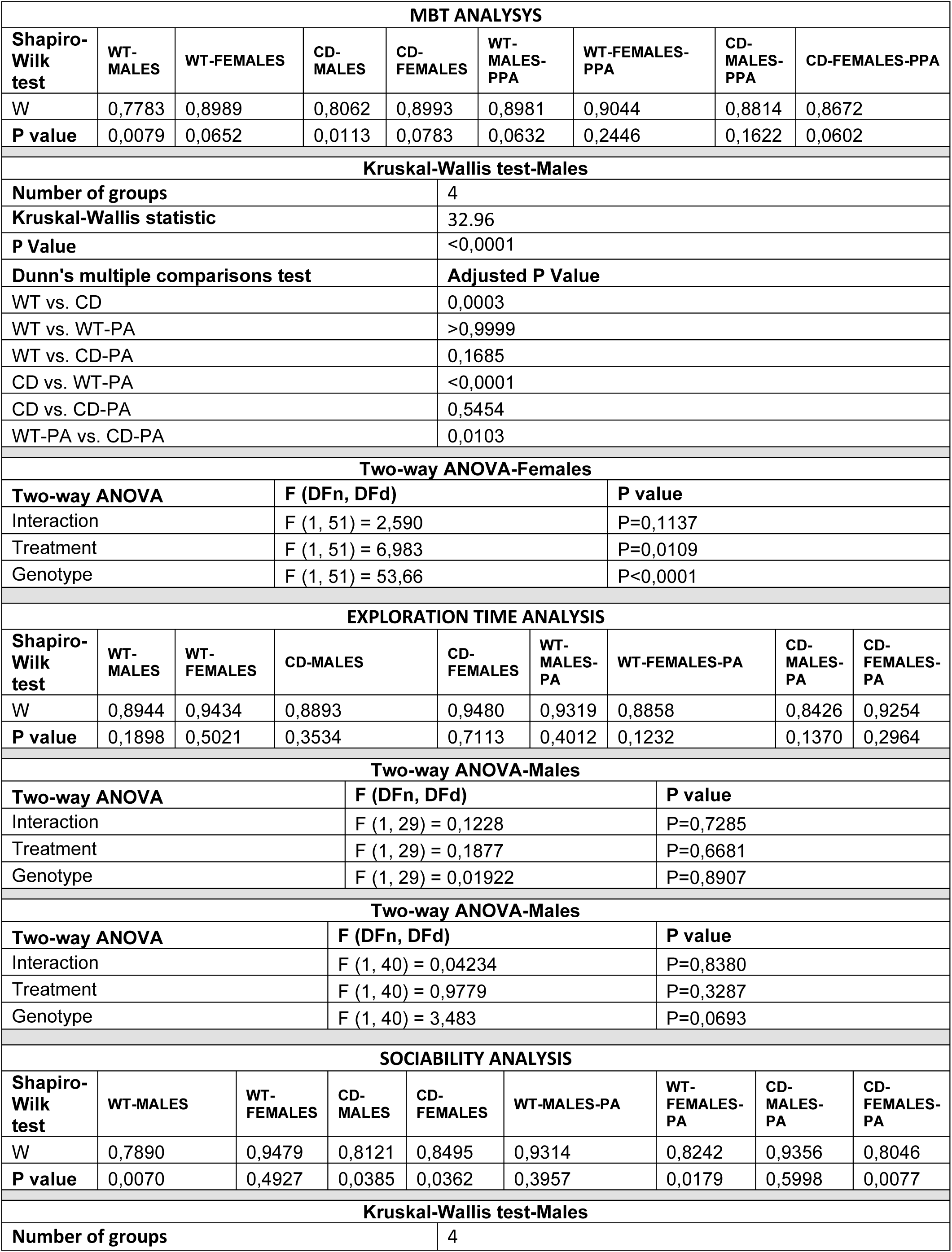
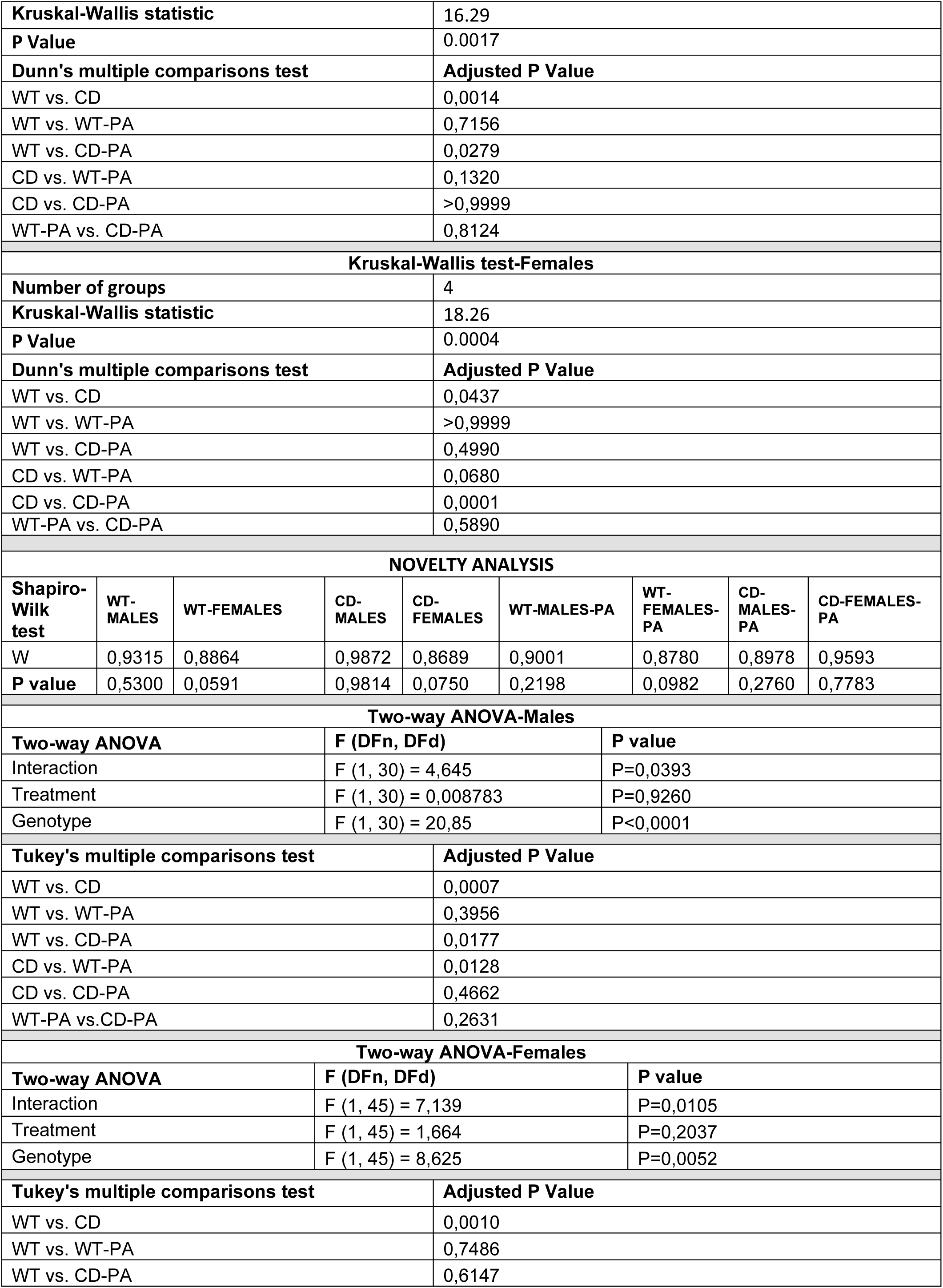
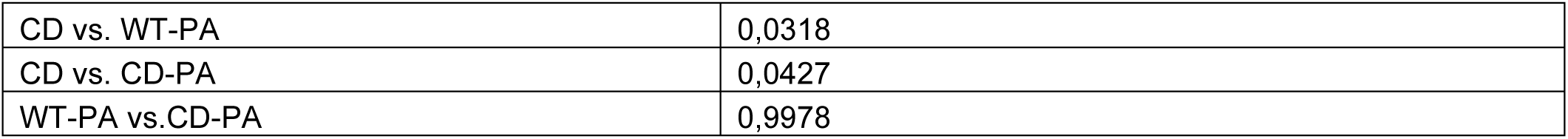
BEHAVIURAL ANALYSYS.

| MBT ANALYSYS |  |  |  |  |  |  |  |  |
| --- | --- | --- | --- | --- | --- | --- | --- | --- |
| Shapiro-Wilk test | WT-MALES | WT-FEMALES | CD-MALES | CD-FEMALES | WT-MALES-PPA | WT-FEMALES-PPA | CD-MALES-PPA | CD-FEMALES-PPA |
| W | 0,7783 | 0,8989 | 0,8062 | 0,8993 | 0,8981 | 0,9044 | 0,8814 | 0,8672 |
| P value | 0,0079 | 0,0652 | 0,0113 | 0,0783 | 0,0632 | 0,2446 | 0,1622 | 0,0602 |
| Kruskal-Wallis test-Males |  |  |  |  |  |  |  |  |
| Number of groups |  |  |  | 4 |  |  |  |  |
| Kruskal-Wallis statistic |  |  |  | 32.96 |  |  |  |  |
| P Value |  |  |  | <0,0001 |  |  |  |  |
| Dunn's multiple comparisons test |  |  |  | Adjusted P Value |  |  |  |  |
| WT vs. CD |  |  |  | 0,0003 |  |  |  |  |
| WT vs. WT-PA |  |  |  | >0,9999 |  |  |  |  |
| WT vs. CD-PA |  |  |  | 0,1685 |  |  |  |  |
| CD vs. WT-PA |  |  |  | <0,0001 |  |  |  |  |
| CD vs. CD-PA |  |  |  | 0,5454 |  |  |  |  |
| WT-PA vs. CD-PA |  |  |  | 0,0103 |  |  |  |  |
| Two-way ANOVA-Females |  |  |  |  |  |  |  |  |
| Two-way ANOVA |  | F (DFn, DFd) |  |  | P value |  |  |  |
| Interaction |  | F (1, 51) = 2,590 |  |  | P=0,1137 |  |  |  |
| Treatment |  | F (1, 51) = 6,983 |  |  | P=0,0109 |  |  |  |
| Genotype |  | F (1, 51) = 53,66 |  |  | P<0,0001 |  |  |  |
| EXPLORATION TIME ANALYSIS |  |  |  |  |  |  |  |  |
| Shapiro-Wilk test | WT-MALES | WT-FEMALES | CD-MALES | CD-FEMALES | WT-MALES-PA | WT-FEMALES-PA | CD-MALES-PA | CD-FEMALES-PA |
| W | 0,8944 | 0,9434 | 0,8893 | 0,9480 | 0,9319 | 0,8858 | 0,8426 | 0,9254 |
| P value | 0,1898 | 0,5021 | 0,3534 | 0,7113 | 0,4012 | 0,1232 | 0,1370 | 0,2964 |
| Two-way ANOVA-Males |  |  |  |  |  |  |  |  |
| Two-way ANOVA |  | F (DFn, DFd) |  |  | P value |  |  |  |
| Interaction |  | F (1, 29) = 0,1228 |  |  | P=0,7285 |  |  |  |
| Treatment |  | F (1, 29) = 0,1877 |  |  | P=0,6681 |  |  |  |
| Genotype |  | F (1, 29) = 0,01922 |  |  | P=0,8907 |  |  |  |
| Two-way ANOVA-Males |  |  |  |  |  |  |  |  |
| Two-way ANOVA |  | F (DFn, DFd) |  |  | P value |  |  |  |
| Interaction |  | F (1, 40) = 0,04234 |  |  | P=0,8380 |  |  |  |
| Treatment |  | F (1, 40) = 0,9779 |  |  | P=0,3287 |  |  |  |
| Genotype |  | F (1, 40) = 3,483 |  |  | P=0,0693 |  |  |  |
| SOCIABILITY ANALYSIS |  |  |  |  |  |  |  |  |
| Shapiro-Wilk test | WT-MALES | WT-FEMALES | CD-MALES | CD-FEMALES | WT-MALES-PA | WT-FEMALES-PA | CD-MALES-PA | CD-FEMALES-PA |
| W | 0,7890 | 0,9479 | 0,8121 | 0,8495 | 0,9314 | 0,8242 | 0,9356 | 0,8046 |
| P value | 0,0070 | 0,4927 | 0,0385 | 0,0362 | 0,3957 | 0,0179 | 0,5998 | 0,0077 |
| Kruskal-Wallis test-Males |  |  |  |  |  |  |  |  |
| Number of groups |  |  |  | 4 |  |  |  |  |
| Kruskal-Wallis statistic |  |  |  |  | 16.29 |  |  |  |
| P Value |  |  |  |  | 0.0017 |  |  |  |
| Dunn's multiple comparisons test |  |  |  |  | Adjusted P Value |  |  |  |
| WT vs. CD |  |  |  |  | 0,0014 |  |  |  |
| WT vs. WT-PA |  |  |  |  | 0,7156 |  |  |  |
| WT vs. CD-PA |  |  |  |  | 0,0279 |  |  |  |
| CD vs. WT-PA |  |  |  |  | 0,1320 |  |  |  |
| CD vs. CD-PA |  |  |  |  | >0,9999 |  |  |  |
| WT-PA vs. CD-PA |  |  |  |  | 0,8124 |  |  |  |
| Kruskal-Wallis test-Females |  |  |  |  |  |  |  |  |
| Number of groups |  |  |  |  | 4 |  |  |  |
| Kruskal-Wallis statistic |  |  |  |  | 18.26 |  |  |  |
| P Value |  |  |  |  | 0.0004 |  |  |  |
| Dunn's multiple comparisons test |  |  |  |  | Adjusted P Value |  |  |  |
| WT vs. CD |  |  |  |  | 0,0437 |  |  |  |
| WT vs. WT-PA |  |  |  |  | >0,9999 |  |  |  |
| WT vs. CD-PA |  |  |  |  | 0,4990 |  |  |  |
| CD vs. WT-PA |  |  |  |  | 0,0680 |  |  |  |
| CD vs. CD-PA |  |  |  |  | 0,0001 |  |  |  |
| WT-PA vs. CD-PA |  |  |  |  | 0,5890 |  |  |  |
| NOVELTY ANALYSIS |  |  |  |  |  |  |  |  |
| Shapiro-Wilk test | WT-MALES | WT-FEMALES | CD-MALES | CD-FEMALES | WT-MALES-PA | WT-FEMALES-PA | CD-MALES-PA | CD-FEMALES-PA |
| W | 0,9315 | 0,8864 | 0,9872 | 0,8689 | 0,9001 | 0,8780 | 0,8978 | 0,9593 |
| P value | 0,5300 | 0,0591 | 0,9814 | 0,0750 | 0,2198 | 0,0982 | 0,2760 | 0,7783 |
| Two-way ANOVA-Males |  |  |  |  |  |  |  |  |
| Two-way ANOVA |  |  | F (DFn, DFd) |  |  | P value |  |  |
| Interaction |  |  | F (1, 30) = 4,645 |  |  | P=0,0393 |  |  |
| Treatment |  |  | F (1, 30) = 0,008783 |  |  | P=0,9260 |  |  |
| Genotype |  |  | F (1, 30) = 20,85 |  |  | P<0,0001 |  |  |
| Tukey's multiple comparisons test |  |  |  |  | Adjusted P Value |  |  |  |
| WT vs. CD |  |  |  |  | 0,0007 |  |  |  |
| WT vs. WT-PA |  |  |  |  | 0,3956 |  |  |  |
| WT vs. CD-PA |  |  |  |  | 0,0177 |  |  |  |
| CD vs. WT-PA |  |  |  |  | 0,0128 |  |  |  |
| CD vs. CD-PA |  |  |  |  | 0,4662 |  |  |  |
| WT-PA vs.CD-PA |  |  |  |  | 0,2631 |  |  |  |
| Two-way ANOVA-Females |  |  |  |  |  |  |  |  |
| Two-way ANOVA |  |  | F (DFn, DFd) |  |  | P value |  |  |
| Interaction |  |  | F (1, 45) = 7,139 |  |  | P=0,0105 |  |  |
| Treatment |  |  | F (1, 45) = 1,664 |  |  | P=0,2037 |  |  |
| Genotype |  |  | F (1, 45) = 8,625 |  |  | P=0,0052 |  |  |
| Tukey's multiple comparisons test |  |  |  |  | Adjusted P Value |  |  |  |
| WT vs. CD |  |  |  |  | 0,0010 |  |  |  |
| WT vs. WT-PA |  |  |  |  | 0,7486 |  |  |  |
| WT vs. CD-PA |  |  |  |  | 0,6147 |  |  |  |

|  |  |
| --- | --- |
| CD vs. WT-PA | 0,0318 |
| CD vs. CD-PA | 0,0427 |
| WT-PA vs.CD-PA | 0,9978 |

**TABLE S8:** XOR BRAIN EXPRESSION.

| XOR BRAIN EXPRESSION-Males |  |  |  |  |  |  |  |  |  |  |
| --- | --- | --- | --- | --- | --- | --- | --- | --- | --- | --- |
| Shapir o-Wilk test | WT- HPP | CD- HPP | WT- TH | CD-TH | WT- SSp | CD- SSp | WT- Ami g | CD- Amig | WT- MOs -p | CD- MO s-p |
| W | 0,9558 | 0,9400 | 0,8087 | 0,9904 | 0,9785 | 0,9751 | 0,8516 | 0,9663 | 0,9705 | 0,9645 |
| P value | 0,7525 | 0,6541 | 0,1188 | 0,9594 | 0,8933 | 0,8728 | 0,2313 | 0,8184 | 0,8444 | 0,8070 |
| MULTIPLE-T TEST |  |  |  |  |  |  |  |  |  |  |
| Brain Region |  | P value |  | t ratio |  | df |  | q value |  |  |
| HPP |  | 0,613421 |  | 0,5326 |  | 6,000 |  | 0,774444 |  |  |
| TH |  | 0,967546 |  | 0,04241 |  | 6,000 |  | 0,977222 |  |  |
| SSp |  | 0,054613 |  | 2,382 |  | 6,000 |  | 0,275794 |  |  |
| Amigdala |  | 0,369046 |  | 0,9710 |  | 6,000 |  | 0,774444 |  |  |
| MOp-MOs |  | 0,583274 |  | 0,5796 |  | 6,000 |  | 0,774444 |  |  |
| XOR BRAIN EXPRESSION-Females |  |  |  |  |  |  |  |  |  |  |
| Sha piro-Wilk test | WT- HPP | CD- HPP | WT- TH | CD-TH | WT- SSp | CD- SSp | WT- Ami g | CD- Amig | WT- MOs -p | CD- MOs-p |
| W | 0,8926 | 0,9207 | 0,9137 | 0,9241 | 0,9287 | 0,9750 | 0,9444 | 0,9098 | 0,8987 | 0,9508 |
| P valu e | 0,3950 | 0,5411 | 0,5022 | 0,5600 | 0,5868 | 0,8722 | 0,6813 | 0,4813 | 0,4244 | 0,7209 |
| MULTIPLE-T TEST |  |  |  |  |  |  |  |  |  |  |
| Brain Region |  | P value |  | t ratio |  | df |  | q value |  |  |
| HPP |  | 0,001039 |  | 5,916 |  | 6,000 |  | 0,004196 |  |  |
| TH |  | 0,007945 |  | 3,904 |  | 6,000 |  | 0,016049 |  |  |
| SSp |  | 0,027106 |  | 2,906 |  | 6,000 |  | 0,036503 |  |  |
| Amigdala |  | 0,212565 |  | 1,395 |  | 6,000 |  | 0,214691 |  |  |
| MOp-MOs |  | 0,686362 |  | 0,4240 |  | 6,000 |  | 0,554581 |  |  |

## Notes

### Competing Interest Statement

The authors have declared no competing interest.

